# Slow-varying normalization explains diverse temporal frequency masking interactions across scales in the macaque visual cortex

**DOI:** 10.1101/2025.03.11.642541

**Authors:** Divya Gulati, Supratim Ray

**Affiliations:** Centre for Neuroscience, Indian Institute of Science, Bangalore, India, 560012

**Keywords:** SSVEP, plaids, low-pass filter, V1

## Abstract

Neurons in the primary visual cortex (V1) respond non-linearly with the presentation of multiple stimuli, which has been explained by a normalization model where the excitatory drive is divided by the summed activity of a large neuronal population. While recent studies have suggested that normalization could be time and frequency-dependent, neural mechanisms underlying this dependence remain unknown. Steady-state visually evoked potentials (SSVEPs), which are produced by presenting flickering or counterphasing visual stimuli, serve as a robust tool to probe these underlying mechanisms by leveraging frequency-specific tagging of concurrently presented stimuli. We presented two overlapping counterphasing grating stimuli (plaids), either parallelly or orthogonally, at multiple contrasts and temporal frequencies and recorded spikes, local field potential, and electrocorticogram from V1 of bonnet macaques while they passively fixated. We also recorded electroencephalogram (EEG) activity. The resulting SSVEPs exhibited complicated dynamics – with “low-pass” and “band-pass” suppression profiles for orthogonal and parallel plaids, respectively. Importantly, these dynamics were conserved across scales – from spiking activity to EEG. Surprisingly, adding a simple low-pass filter in the normalization signal sufficiently explained these diverse effects. Our results present a simple mechanism to explain the spectro-temporal dynamics of normalization. These insights may also aid in designing and interpreting SSVEP-based cognitive and EEG-based brain-computer interfacing (EEG-BCI) studies.

## INTRODUCTION

For efficient encoding of sensory information, neurons dynamically adjust their responsiveness based on the context^1,2^. In addition, visual cortical neurons often exhibit non-linear response properties when multiple stimuli are presented, such as cross-orientation suppression^3,4^, surround suppression^5^, and pattern masking^3,6–11^. Many of these response properties can be explained using a canonical computation known as the divisive normalization model, which asserts that the pooled activity of the neighbouring neuronal population scales a given neuron’s excitatory drive^12^. The model can also be generalized to various sensory modalities, cognitive functions, and species^13–15^. However, the basic normalization model does not consider the dynamic temporal facets of the neural responses. Though a growing body of literature^6,16–19^ suggests that the addition of a slowly accumulating inhibitory drive can explain changes in the temporal domain, the factors influencing the summation of signals within the inhibitory pool are not yet fully teased apart, especially for temporally modulated inputs.

These underlying rules can be probed using masking paradigms, which involve simultaneously presenting multiple stimuli in spatial proximity^20^. Temporal frequency-based masking interactions can be studied by presenting a plaid stimulus generated by overlapping two counterphase or contrast reversal gratings. A counterphase stimulus evokes a stimulus-locked periodic oscillatory response at the temporal frequency of the stimulus^21^, known as steady-state visual evoked potential (SSVEP). SSVEPs are very sensitive to changes in stimulus characteristics, have a high signal-to-noise ratio, and are resistant to artefacts^22,23^, which makes them suitable for cognitive research paradigms and EEG-based BCI studies^23–25^. Two stimuli tagged at different temporal frequencies generate SSVEPs at the respective stimulus frequencies, their harmonics, as well as low-order sums and differences of the input frequencies^26^. Therefore, SSVEPs could also be used to quantify the suppression in the response of a stimulus (referred to as ‘target’) when a second stimulus (‘mask’) is presented. Normalization-based models have been used to understand the SSVEP and its harmonic responses^6,8,9^.

Figure 1A shows a normalization model along with the equation^8,27^. Generally, the target grating is kept at a fixed temporal frequency while the temporal frequency of the mask is varied. The response is considered only at the target frequency. The excitatory drive only depends on the contrast of the target frequency and has no contribution from the mask stimulus since the mask has a different temporal frequency. Nevertheless, non-linear interactions allow the mask to contribute to the normalization pool (the term S in the denominator of the equation in Figure 1A denotes this contribution). This suppressive contribution (S) could further depend on the difference between the orientations of the target and mask (δθ) or the difference in the target and mask frequency (δ*f*).

**Figure 1.**
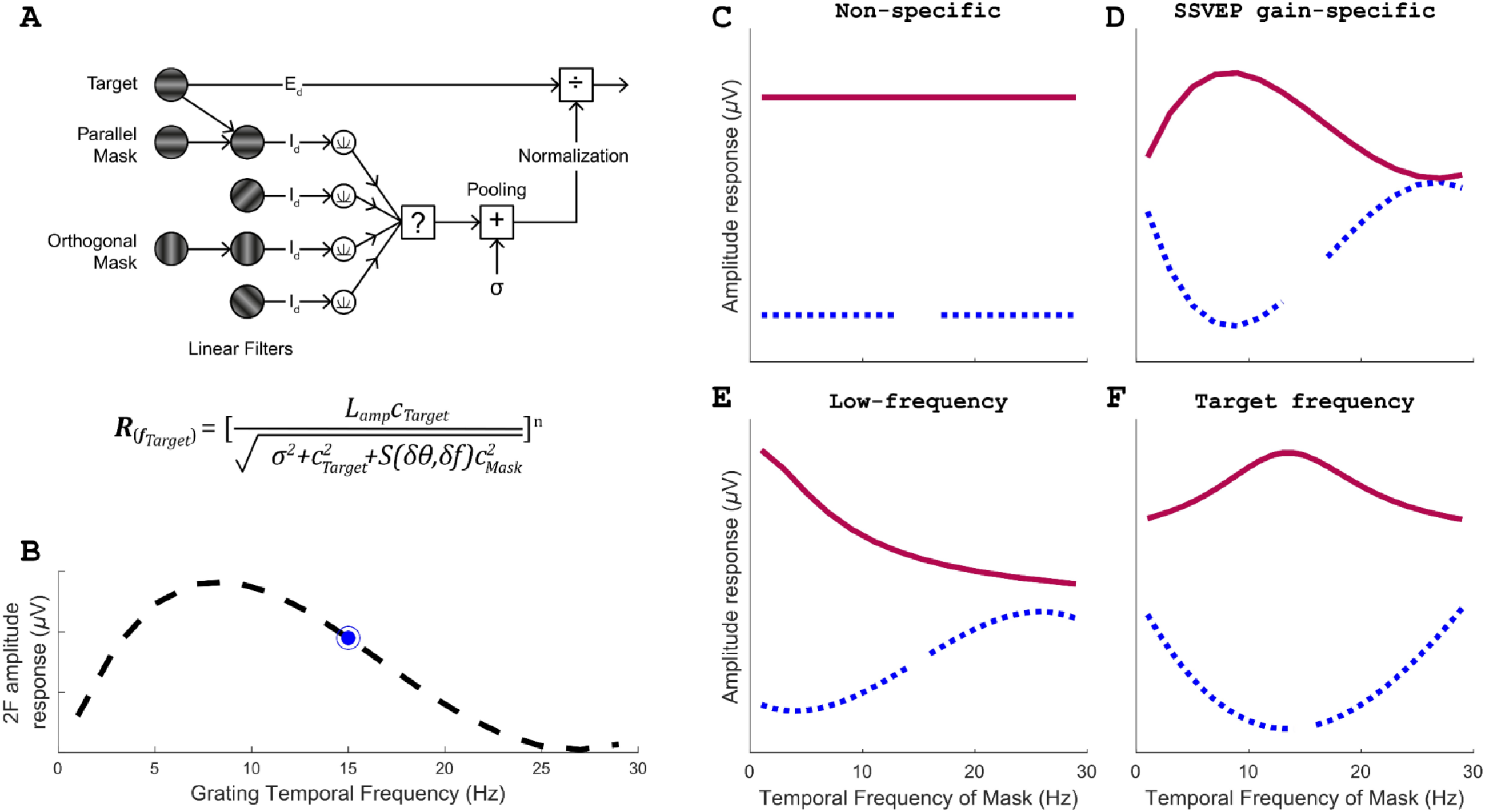
Hypothesis for interaction between target and mask temporal frequency. (A) Schematic and equation for the tuned normalization model. The masking model is adapted from Foley’s Model 3^8,27^. The excitation drive only consists of input from the target grating. The inhibitory drive consists of input both from the target and mask grating. The filters tile the entire orientation space (from 0° to 180°). For the parallel mask, the two gratings match in their orientation and, thus, pass through the same filter. They are, therefore, first summed and then passed through expansive non-linearity. For an orthogonal mask, the two pass through different filters and are passed through expansive non-linearity separately and then summed. This drive is then pooled, and it divisively scales the excitatory drive. However, the temporal dynamics of pooling are not well characterized (denoted by the question mark). (B) SSVEP tuning profile of counterphase grating presented at 25% contrast. The blue dot represents the target frequency used in the main study. (C-F) Hypothetical suppression profile (solid deep magenta trace) as a function of temporal frequency of mask grating, and the corresponding response profile (dotted blue trace) at the temporal frequency of the target grating. (C) Non-specific suppression: Suppression strength is independent of the temporal frequency of the mask. (D) SSVEP gain-specific suppression: Suppression strength is dependent on SSVEP tuning preference as shown in B. (E) Low-frequency tuned suppression: Suppression strength is progressively stronger for lower frequencies. (F) Target frequency dependent suppression: Suppression strength is maximum when the temporal frequency of the mask grating is closer to the frequency of the target grating and decreases as the difference between the temporal frequencies of the two gratings increases.

There could be several hypotheses as to how the response of the target grating could vary as a function of the temporal frequency of the mask (Figure 1, panel C-F; suppression profile (S) is shown in magenta, while the response profile is shown in dotted blue). Suppression could be non-specific (Figure 1C) or could be proportional to the magnitude of the SSVEP response at different frequencies (Figure 1D; typical SSVEP response profile in primate primary visual cortex (V1) is shown in Figure 1B). Other possibilities include having stronger suppression from low-frequency masks (Figure 1E) or target-frequency dependent suppression that is maximally suppressive at mask frequencies closest to the target frequency (Figure 1F). These hypotheses can be tested by presenting masks and targets at multiple frequencies. While we initially found evidence for low-frequency suppression for parallel plaids in the local field potential (LFP) recorded from V1 of monkeys^9^, the profile was found to be target-frequency dependent (Figure 1F) when a larger range of target and mask frequencies were used, in both LFP and human EEG recordings^10^. For orthogonal plaids, the profile was non-specific (Figure 1C).

However, three questions remain unanswered. First, these studies used full-field stimuli that did not elicit robust spiking activity, so it is unclear if spikes also show similar response characteristics as LFP. Second, previous studies used a single mask contrast, so the suppression profile was not explicitly estimated (only the response profiles were compared). Most importantly, there was no explanation for the hypothesized suppression profiles. To address these questions, we presented small plaid stimuli at multiple target and mask contrasts while recording chronically from macaque V1 and directly estimated the suppression profiles for both LFP and spiking responses for both parallel and orthogonal plaids. Extending previous results, we found target-specific (Figure 1F) and low-frequency (Figure 1E) suppression profiles for parallel and orthogonal plaids across all scales – spikes, LFP. ECoG and EEG. Surprisingly, simply adding a low-pass filter in the inhibitory drive (in the box marked with “?” in Figure 1A) was sufficient to explain these varied effects.

## RESULTS

We trained four monkeys to passively fixate at a fixation point within a 2° window. At the same time, we presented either a full-field or a small (1° or 1.5°) counterphase grating or plaid stimulus centered on the receptive fields (eccentricity of ∼2.5°-4.5°; Figure S1) of the microelectrode grid. We recorded spiking and LFP activity using a chronic array from two monkeys (M1 and M2) and ECoG activity using a custom hybrid array from a third monkey (M3), and EEG activity from two monkeys (M2 and M4) while they performed the fixation task. SSVEPs generated in response to counterphase gratings or plaids were characterized using their amplitude spectrum computed using Fourier analysis. In cases of plaid stimuli, the target grating had a fixed temporal frequency of 15 Hz across all experiments, and the mask grating had a varying temporal frequency between 1 and 29 Hz across trials.

### Response profiles remain similar across stimulus sizes and recording scales

First, we tested whether the response profiles reported earlier for full-field stimuli^9,10^ could be observed with small stimuli. The mask grating was fixed at 25% contrast, whereas the target grating could be either at 0 or 25%. This allowed us to have both plaid and grating conditions in the same experiment. The relative orientation difference between the two gratings was either kept at 0° or 90°, giving parallel and orthogonal plaids, respectively. Figure 2, panels A and B, show the amplitude spectrum of electrode-averaged LFP responses for target grating only (grey trace; same for all conditions) and parallel and orthogonal plaid conditions (colour traces) for full-field and small stimuli, respectively. These counterphasing stimuli produce the strongest SSVEP response at twice the stimulus frequency^9,10^, so we analyzed the change in amplitude for the target grating at 30Hz. The response at 30Hz for plaid conditions was reduced compared to the grating condition when the mask temporal frequency differed from that of the target grating (shown by red bars). Facilitation only happened when the target and mask had the same temporal frequency. The suppression in response to a plaid stimulus was quantified as the change in amplitude between grating-only and plaid conditions. We found that the response profiles for full-field (Figure 2C) and small (Figure 2D) were similar, with both showing a target frequency-dependent profile (Figure 1F) for parallel and non-specific (Figure 1C) profile for orthogonal plaids, as shown previously for full-field stimulus^9,10^. Suppression was stronger for parallel plaids than orthogonal, as observed previously in macaques and humans^8–10,27^. In addition to self-harmonics peaks in the amplitude spectra, we observed peaks at intermodulation (IM) terms (*f_1_+f_2_, f_1_-f_2_, 2f_1_+2f_2_, 2f_1_-2f_2_*) primarily for parallel plaid conditions.

**Figure 2.**
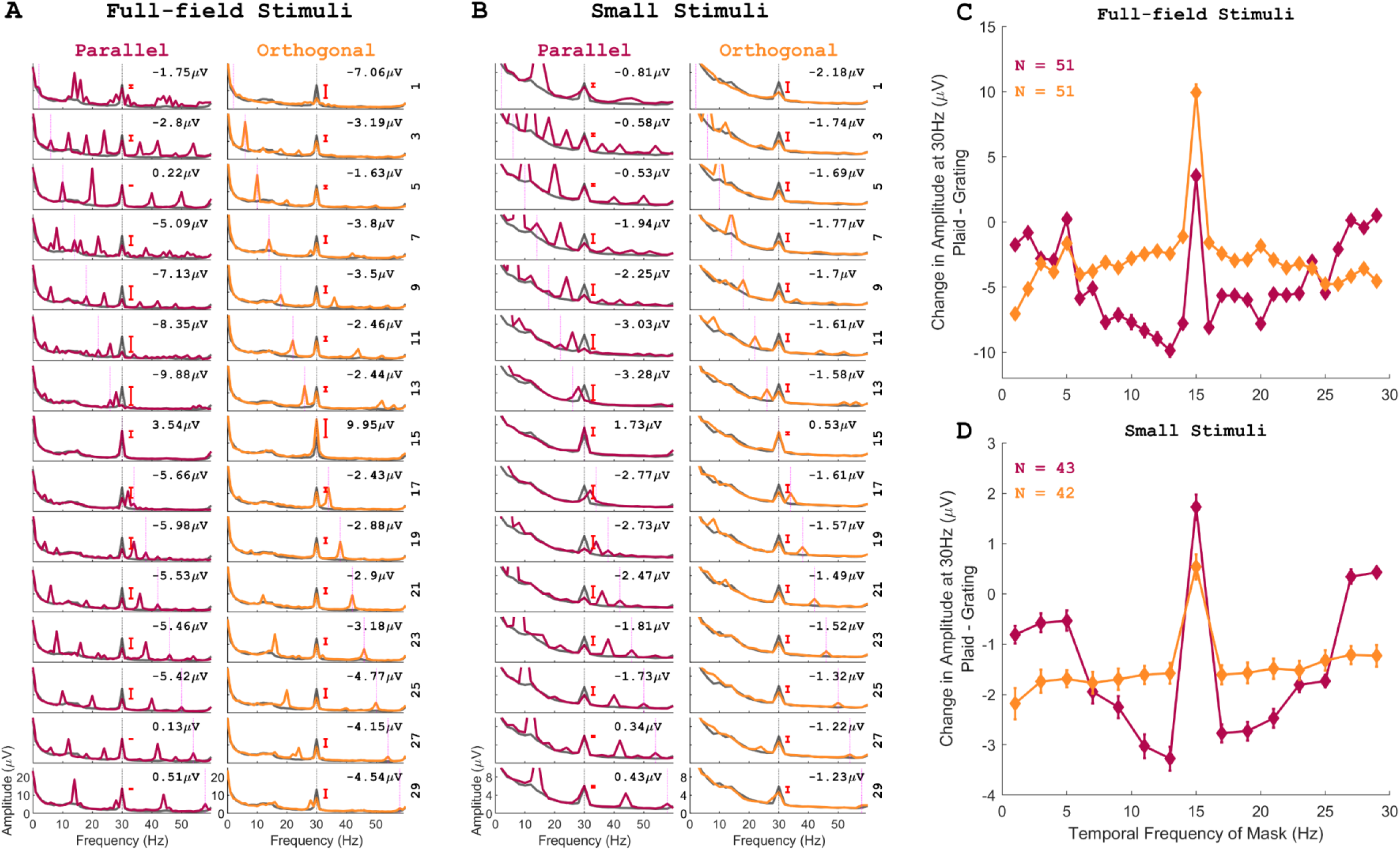
LFP amplitude response suppression for different stimuli sizes. (A) Mean amplitude spectra averaged across M1 and M2 microelectrodes for a full-field stimulus. (B) Same as (A) but for a small stimulus presented over the aggregate receptive field of the microelectrode grid. The two columns in (A) and (B) indicate the two different plaid stimuli (deep magenta – parallel; gold – orthogonal), indicating the relative orientation difference between the target and mask grating, and each row corresponds to a different temporal frequency of mask grating superimposed on the target grating of 15Hz (indicated on the right side of each plot). The grey coloured trace denotes the grating ‘only’ condition at 25% contrast (same across all the plots for (A) and across each column for (B) as a single type of plaid stimuli was presented in one session for small-sized stimulus). The coloured trace represents the plaid condition with target and mask grating at 25% contrast each. The dotted vertical lines indicate the second harmonic of the target (black) and mask grating (magenta). The numbers indicate the difference in amplitude between the grating ‘only’ condition and the relative plaid condition at 30Hz. (C) Amplitude response difference between the full-field grating ‘only’ condition and plaid condition as a function of the temporal frequency of the mask grating, averaged across electrodes for two monkeys (N = 51). Error bars indicate ±SEM. (D) Same as (C) but for small stimuli (averaged across electrodes for all sessions for parallel (N = 43) and orthogonal plaids (N = 42)).

While these stimuli allow us to obtain the response profiles (Figure 2C-D), the suppression profile S(δθ,δ*f*) in Figure 1A cannot be determined from this data. This is because S(δθ,δ*f*) has a different value for each mask temporal frequency (14 free parameters each for parallel and orthogonal conditions in our case). Therefore, to estimate S(δθ,δ*f*) for a particular mask frequency and orientation, multiple mask and target contrasts are needed, which was not done in our previous studies. To address this, we presented both target and mask gratings independently at 4 contrasts (0%,6.25%,12.5% and 25%), generating 16 response profiles (one for each combination of target and mask contrasts; Figure 3).

**Figure 3.**
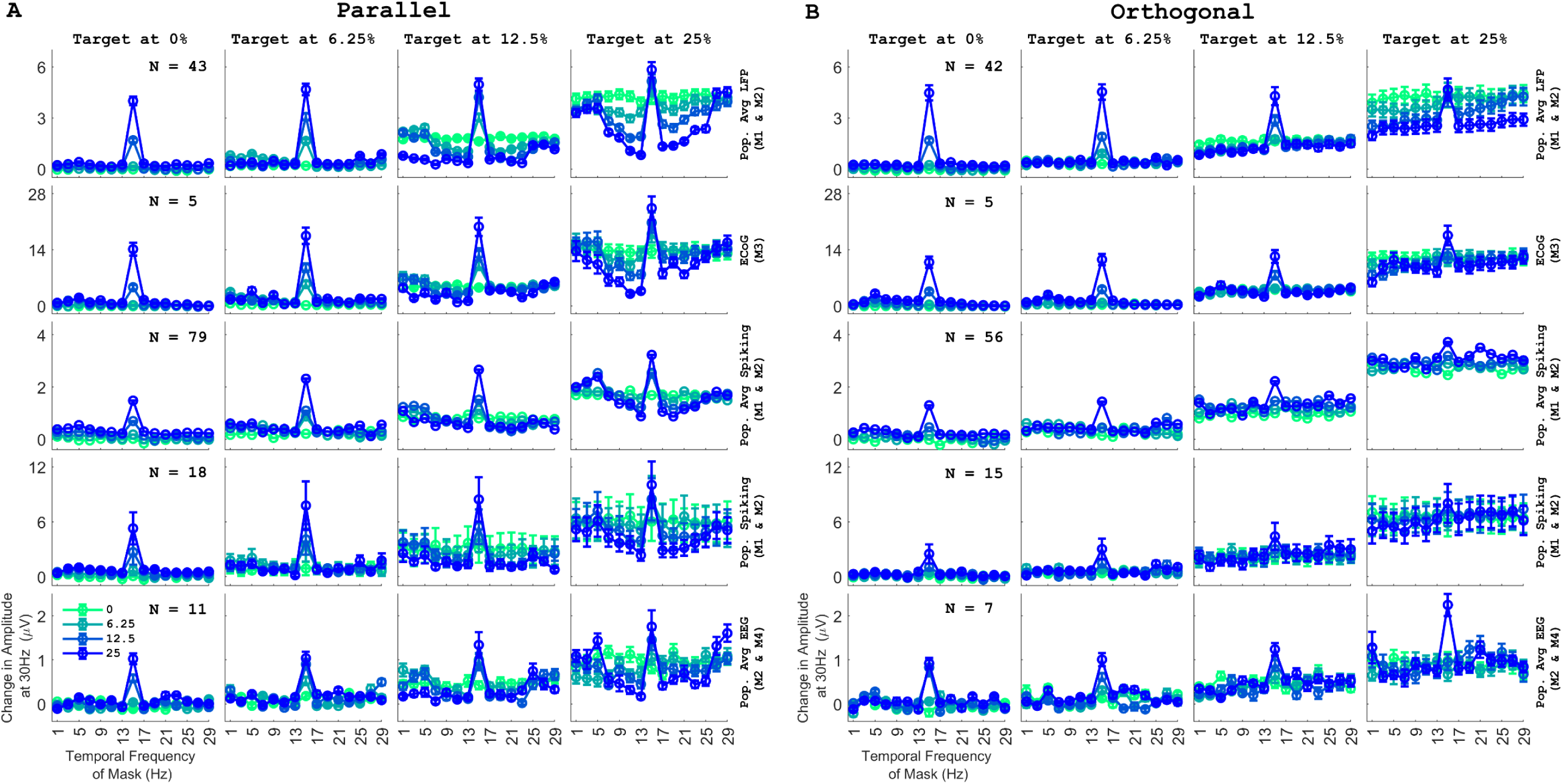
Amplitude response suppression as a function of different contrasts and orientations across recording scales. (A) Change in amplitude at 30Hz from the baseline condition averaged across sessions and electrodes for parallel plaid stimuli, with target and mask grating presented at various contrast combinations plotted as a function of temporal frequency of the mask grating. Each grating can take one of the four contrast values between 0% and 25% (on a log scale). Within each panel, four traces represent the contrast of the mask grating (green to blue, indicating increasing contrast level), and each column represents the contrast of the target grating. Different row corresponds to different recording scales. The first row represents the average change in amplitude response from the baseline condition for M1 and M2 for a small stimulus in the LFP signal. The second row corresponds to the change in amplitude from the baseline for the ECoG response in M3 against a full-field stimulus. The third row indicates the average change in amplitude calculated from the Fourier transform of the trial and electrode-averaged peri-stimulus time histogram (PSTH) for M1 and M2 in response to the presentation of a small stimulus. The fourth row represents the change in amplitude calculated from the Fourier transform of the PSTH of selected electrodes from M1 and M2. Only electrodes with high firing rates during the stimulus period were selected. The last row represents the change in amplitude response for EEG activity in M2 and M4 against a full-field stimulus. Error bars indicate ±SEM. ‘N’ indicates the number of electrodes. (B) Same as (A) but for orthogonal plaids.

We first tested whether the SSVEP interactions at the LFP/ECoG/EEG scale differ from spiking activity. We presented stimuli big enough to cover the aggregate receptive fields of the microelectrode grids for LFP and MUA recordings, while full-field stimuli were presented for ECoG recordings (as the receptive field centers were far apart; Figure S1) and also for EEG recordings. Figure 3A-3B shows the response profiles for varying mask and target contrasts for LFP (top row), ECoG (second row), spikes (third and fourth row), and EEG (bottom row). To observe changes at the spiking level, we first generated a single response profile after averaging all the spiking data across electrodes and then computing the amplitude at 30 Hz (third row).

This serves as a proxy for LFP signals, which reflect the aggregate activity of hundreds of neighbouring neurons. Note that the firing rates must be sufficiently high to get a robust response at 30Hz during stimulus presentation using Fourier analysis for an individual electrode. So, we subselected electrodes with high-firing rates during the sustained period (0.25-0.75ms) and calculated the Fourier transform on the PSTH of each electrode (fourth row). Importantly, we found that the characteristic dip in the response profile near the target frequency, characteristic of the target-frequency profile (Figure 1F), was present even for spiking data for mask and target contrasts of 25% (blue trace in the rightmost column of Figure 3A; third and fourth row). Similarly, for orthogonal plaids (Figure 3B), spiking responses showed a non-specific response similar to what was observed at other recording scales. This was observed in EEG signals as well. These results show that the core shape of the response profile is visible even for individual neurons, even though it has been previously observed that spikes, LFP, and EEG have different tuning for preferred temporal frequencies^9^.

In either of the recording scales (spikes/LFP/ECoG or EEG), suppression became stronger with increasing target contrast and mask contrast. The suppression profile appeared contrast invariant for parallel plaids, such that increasing contrast only scaled the responses but did not change the target-frequency-dependent profile. Interestingly, for orthogonal plaids with lower contrasts of mask or target, a low-frequency tuned suppression was observed instead of a non-specific suppression observed when both target and mask were at maximum contrast (25% each). We also recorded responses for plaids with component gratings separated by 45° (Figure S2, panel A) in orientation for M2 and M3. We observed a similar low-frequency tuned suppression in LFP and ECoG activity.

### The suppression profile shape is based on the relative orientations of the component gratings

In masking literature, normalization models have been extensively used to characterize responses to temporally modulated plaid stimuli^3,6–9^ as they capture non-linear operations well. So, we examined if our observations could also be fitted using a ‘tuned’ normalization model^9^ (Figure 1 panel A, equation 4 in <u>methods</u>). This equation has 16 free parameters – L_amp,_ which denotes response within the receptive field to a stimulus at unit contrast, σ (semi-saturation constant), and 14 ‘S’ parameters to describe the suppression profile due to 14 different temporal frequencies of mask grating. We fitted this model to data from individual spiking, LFP, ECoG, and EEG electrodes (Figure 4). For each electrode, the entire dataset consisting of 14 mask temporal frequencies (the data point at the target frequency was not considered) and 16 different contrast combinations (224 data points as shown in individual rows of Figure 4) was fitted using this equation. The predictions (Figure 4) explained more than 80% of the variance for almost all conditions (except for the orthogonal condition in M1, as the recording quality for that session was noisy) in LFP and ECoG signals (>60% variance for multiunit and EEG activity). We obtained two kinds of suppression profiles depending on the relative orientation difference between the component counterphase gratings. We obtained a band-pass-like suppression profile for parallel plaids (Figure 4 panel A, last column, deep magenta trace), and a low-frequency tuned profile for orthogonal and 45 separated plaids (Figure 4 panel B, last column, gold trace; Figure S2 panel B-C, top rows, last column, orange trace). However, the suppression profile for the parallel case had two distinct features – suppression values tended to be higher at very low frequencies (especially pronounced for M1), and there was asymmetry around the target frequency as well, with mask frequencies lower than the target, producing stronger suppression than those higher. This asymmetry is also readily observed in the response profiles for the target contrast of 25% (Figure 4A).

**Figure 4.**
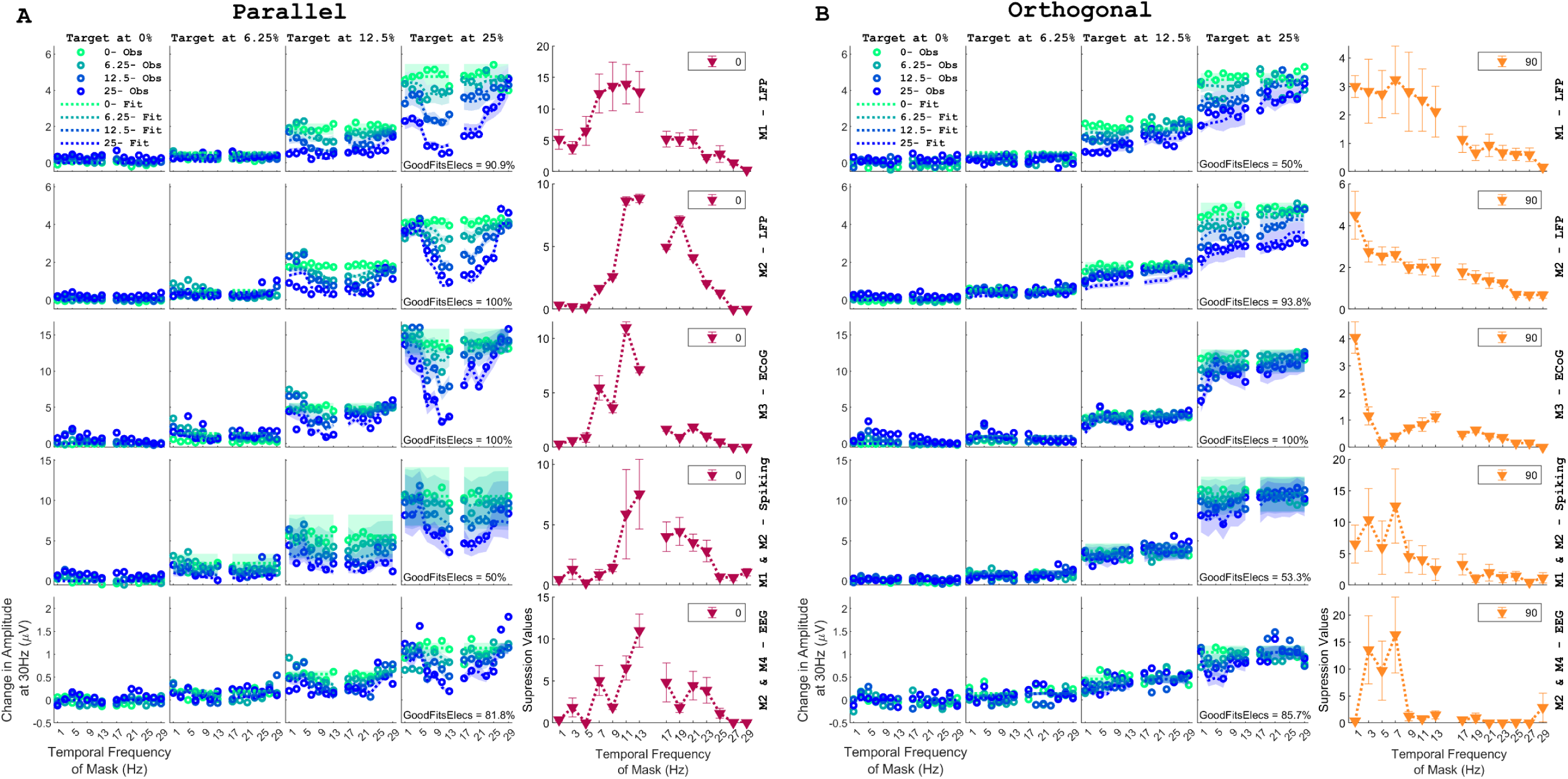
The suppression profile obtained from the tuned normalization model depends on the relative orientation difference between the target and mask grating. (A) Model fits for averaged amplitude response at 30Hz and suppression profiles obtained from the tuned normalization model (equation 4) for parallel plaids. The first three rows correspond to the LFP response for M1 and M2, and the ECoG response for M3; plotted for electrodes with explained variance >= 0.8. The fourth row corresponds to multiunit activity obtained from M1 and M2. The last row represents the EEG activity recorded from M2 and M4. MUA and EEG activity are plotted for electrodes with explained variance >= 0.6. The first four columns show the model fits (dotted traces with the shaded region indicating mean±SEM) for the empirically averaged amplitude response (circles, error bars not shown) for different target contrasts. Each subplot further has four coloured traces corresponding to different mask contrasts. Same as Figure 3. The percentage of electrodes with explained variance >= 0.8 for LFP and ECoG activity, and >=0.6 for multiunit and EEG activity, is represented as ‘GoodFitsElecs’ in the inset of each row. The last column of each row represents the mean suppression values obtained from the model, with error bars indicating ±SEM. (B) Same as (A) but for orthogonal plaids.

To limit the number of free parameters, we fixed the exponent value to 2. However, we also fitted the data by keeping the exponent as a free parameter and found that these exponent values converged near 2 (data not shown). The median values obtained for semi-saturation constant (σ) and L_amp_ are shown in Table 1–3 (for parallel and orthogonal plaids for: 1. LFP and ECoG, 2. for spiking, 3. for EEG) and Table S1 (for 45° separated plaids). The original-tuned normalization model used here does not assume any functional form for the suppressive drive. So, our next goal was to understand how the shape of the suppression profile can be obtained.

**Table 1:** Showing median ± bootstrapped SEM values for parameters obtained from the original-tuned normalization model for LFP and ECoG data.

|  | *Parallel Plaid* |  |  | *Orthogonal Plaid* |  |  |
| --- | --- | --- | --- | --- | --- | --- |
| *Parameters* | M1 | M2 | M3 | M1 | M2 | M3 |
| $\sigma$ | 0.23±0.11 | 0.23±0.01 | 0.31±0.01 | 0.21±0 | 0.23±0.03 | 0.35±0.01 |
| $L_{amp}$ | 3.41±0.57 | 2.58±0.21 | 6.22±0.12 | 2.88±0.06 | 2.45±0.55 | 6.39±0.24 |

**Table 2:** Showing median ± bootstrapped SEM values for parameters obtained from the original-tuned normalization model fitted for spiking data.

|  | *Parallel Plaid* | *Orthogonal Plaid* |
| --- | --- | --- |
| *Parameters* | M1 and M2 | M1 and M2 |
| $\sigma$ | 0.44±0.12 | 0.40±0.13 |
| $L_{amp}$ | 5.49±0.75 | 6.43±0.6 |

**Table 3:**
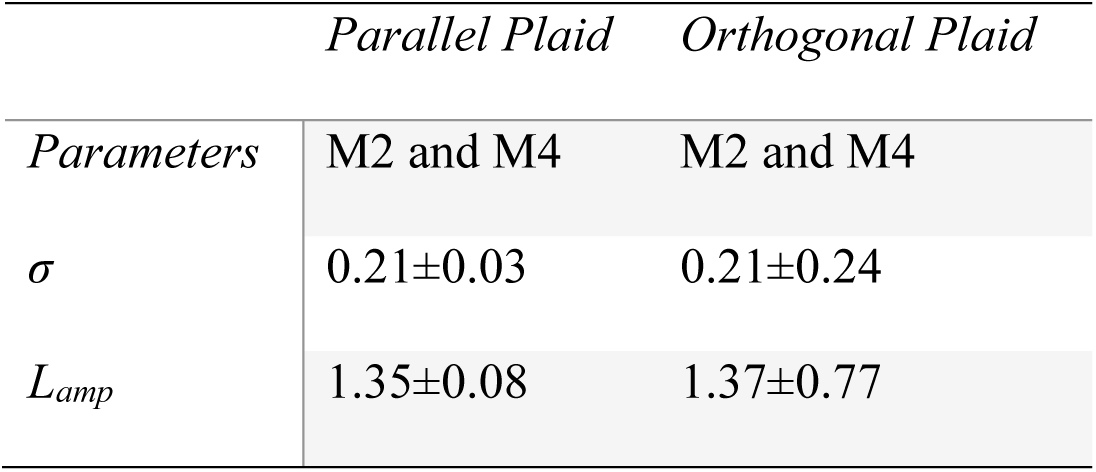
Showing median ± bootstrapped SEM values for parameters obtained from the original-tuned normalization model fitted for EEG data.

### Low-pass filtering of the normalization signal produces the desired suppression profiles

Previous reports^8,27,28^ suggest that the inhibitory drive is a combination of the physical form of the input stimuli. So, we first assumed the ‘S’ as a summation of two sinusoids corresponding to target and mask gratings, and to account for the stronger response at the second harmonics in the response (owing to the squared output of complex cells^29,30^), these signals can be raised to a squaring exponent. For parallel plaids, summation happens before squaring since the signals pass through the same channel (see Figure 1A), while for orthogonal plaids, the sinusoids are squared and then summed. The first column in Figure 5 shows these signals for the fixed target frequency and three different mask frequencies, while the second column shows the normalization signals, with 5A and 5B representing the computation for parallel and orthogonal conditions. The overall normalization strength obtained by averaging this signal is comparable whether the mask frequency is slower than target (pinkish-purple trace), near the target (blue) or faster than target (cyan), and therefore the suppression profile is essentially flat (third column; at the mask frequency of 5 Hz there is an overlap between an IM term leading to a different value in Figure 5A). Thus, the basic normalization model yields a non-specific suppression and response profile.

**Figure 5.**
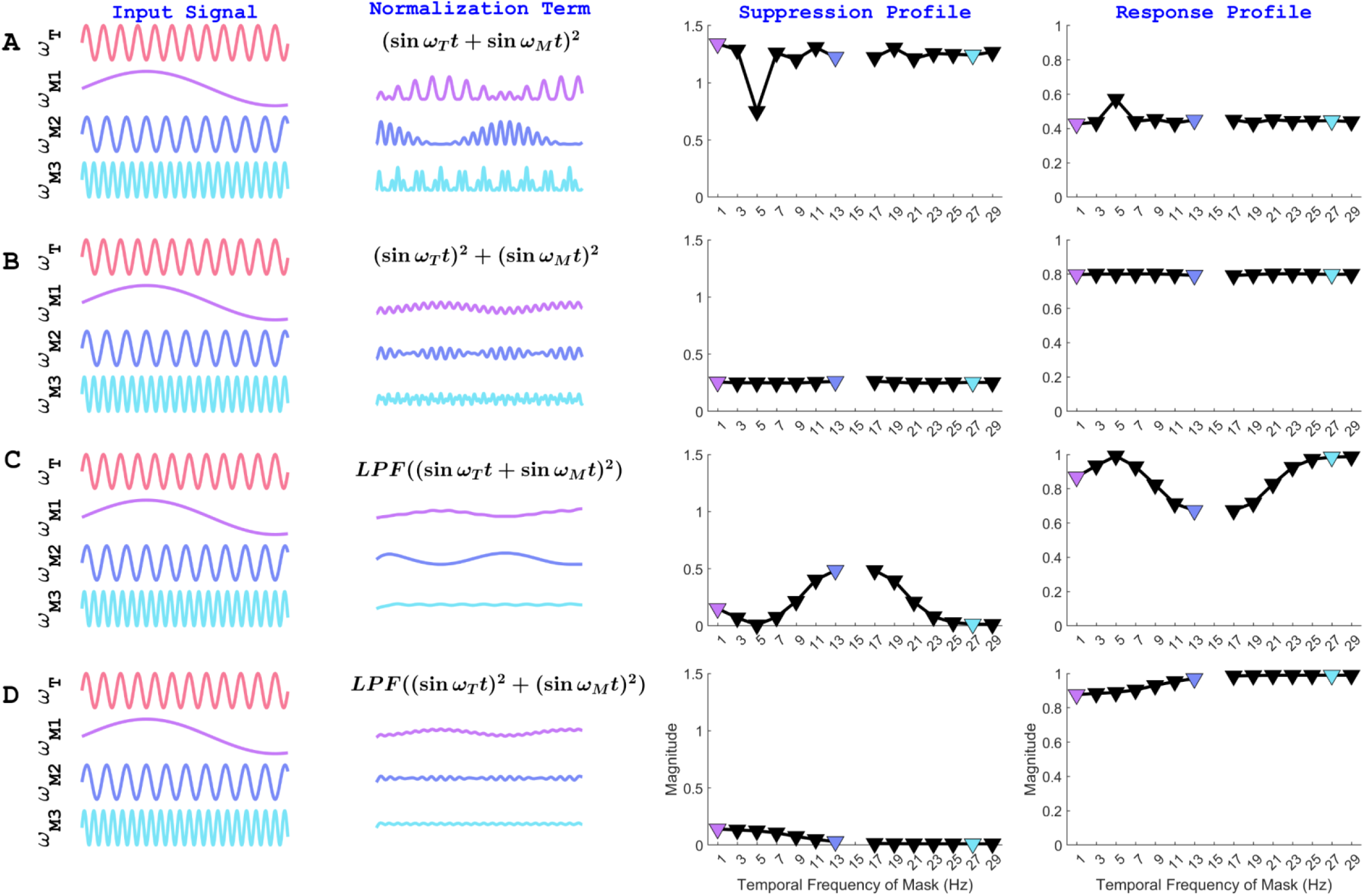
Simulations of the drive coming from the normalization pool and their expected suppression profiles. The four columns correspond to the input signal, inhibitory normalization drive based on different equations, expected suppression profiles, and response profiles. Three different mask gratings (same across A-D) of varying temporal frequencies are highlighted in different colours. The input signal represents the form of the sinusoids. The normalization term depicts the input signal after going through the operation described. The suppression profile is obtained by calculating the energy of the inhibitory drive by squaring the amplitude of the response across the entire spectrum (equation 5). The response profile is the inverse of the magnitude of the suppression profile. (A) The inhibitory normalization signal is obtained by first adding the signal from the target and mask sinusoids, followed by rectification. (B) The normalization signal is obtained by rectifying the target and mask sinusoids and then summing up the input. (C) Same as (A), but the inhibitory drive is passed through a low-pass filter, analogous to integrating responses from the neighbouring population over some temporal integration window. (D) Same as (B), but the inhibitory drive passes through a low-pass filter.

Some studies^6,16–19^ have suggested that the normalization signal is not instantaneous with respect to the incoming excitatory drive leading to response onset. The delay in the signal can be achieved by low-pass filtering the normalization signal. We found that simply adding a low-pass filter (LPF) to the normalization signal shown in the second column is sufficient to get the desired suppression profiles. For parallel plaids, this happens because when two sinusoids are summed and squared, the output contains terms at harmonics, sum and difference intermodulation frequencies:

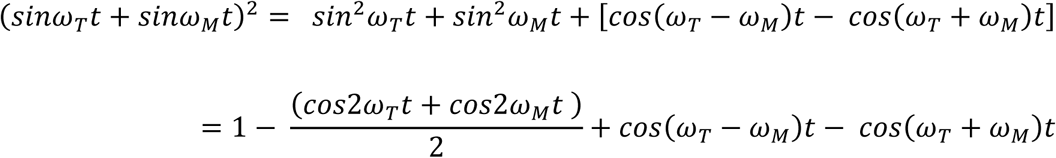

When this signal is passed through a low-pass filter, the output is high when (i) lower mask frequencies (ω_*M*_) are presented or when (ii) ω_*M*_is close to ω_*T*_ such that the sinusoid with frequency of |ω_*T*_ − ω_*M*_| can pass through the filter. Therefore, maximum suppression (Figure 5C, third column) is observed for frequencies closest to the target frequency, followed by slower mask frequencies (Figure 5C, third column). Indeed, the low-pass-filtered normalization signal shown in the second column of Figure 5C is largest when the mask is close to the target frequency (blue trace), followed by the slow mask frequency (pinkish-purple trace), and is weakest for fast mask frequencies (cyan trace). Similarly, the low-frequency dependent suppression profile can be obtained when sinusoids, which are first squared and then summed, are passed through a low-pass filter (Figure 5D). In this case, there are no intermodulation terms, so lower mask frequencies are most suppressive since they can pass through the low-pass-filter while higher mask frequencies cannot (Figure 5D).

In this formulation, there is no explicit suppression profile (S) anymore that is contributed by the mask, since both target and mask frequencies interact non-linearly to produce the overall normalization signal. However, to maintain uniformity with the previous model, we continued with a suppression profile (as in equation 4) which was explicitly calculated by the action of a low-pass filter on the summed and squared (or squared and summed) sinusoids. To account for the effect of orientation, we assumed that the suppression profile was a weighted sum of the two summation mechanisms described in Figure 5C and 5D (equation 5), with the weights as free parameters dependent on the relative orientations of the gratings.

Next, we fitted our data with this ‘optimal-tuned normalization model’, where the number of free parameters to represent S reduced from 14 to 6 (equation 5). Despite that, we could explain >80% of the variance for more than ninety percent of the electrodes (Figure 6) in LFP and ECoG data and could qualitatively capture the overall response and suppression profiles. We obtained a target frequency-dependent profile for the parallel plaids with α_1_ > α_2_ (Table 4) as well as a low-frequency tuned profile for orthogonal plaids with α_2_ > α_1_. In multiunit spiking activity, we observed similar suppression profiles with more than 60% explained variance for about half of the units (Table 5). In EEG activity, we could also capture >60% explained variance for more than 60% of the electrodes with comparable parameter values (Table 6) to those obtained for other recording scales. For plaids with 45° separation (for M2 and M3; Figure S2 panel B and C bottom rows), a low-frequency tuned suppression profile was obtained with more than 80% explained variance for all the electrodes and α_2_ > α_1_ (see Table S2). Our empirical data suggested that distinct channels give rise to orientation-selective masking profiles.

**Figure 6.**
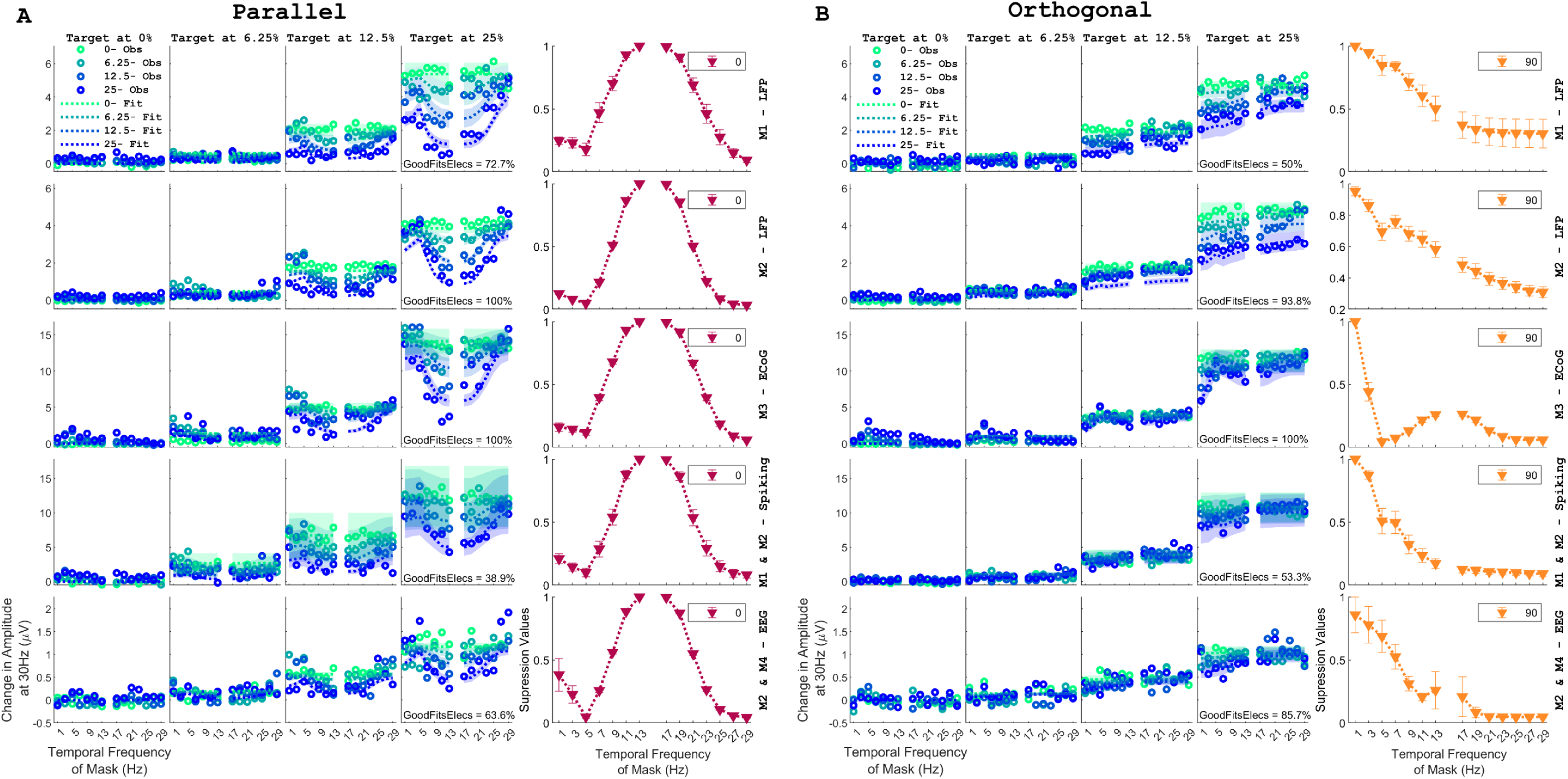
An optimal-tuned normalization model with a low-pass filtered normalization signal can explain SSVEP suppression responses. (A) and (B) same as Figure 4, but the model is fitted using Equation 4 and Equation 5. The suppression profiles as a function of temporal frequency of mask grating (in the last column; shown in magenta and gold) were calculated from the free parameter values obtained from the optimal-tuned normalization model using Equation 5 and were normalized for each electrode before averaging.

**Table 4:**
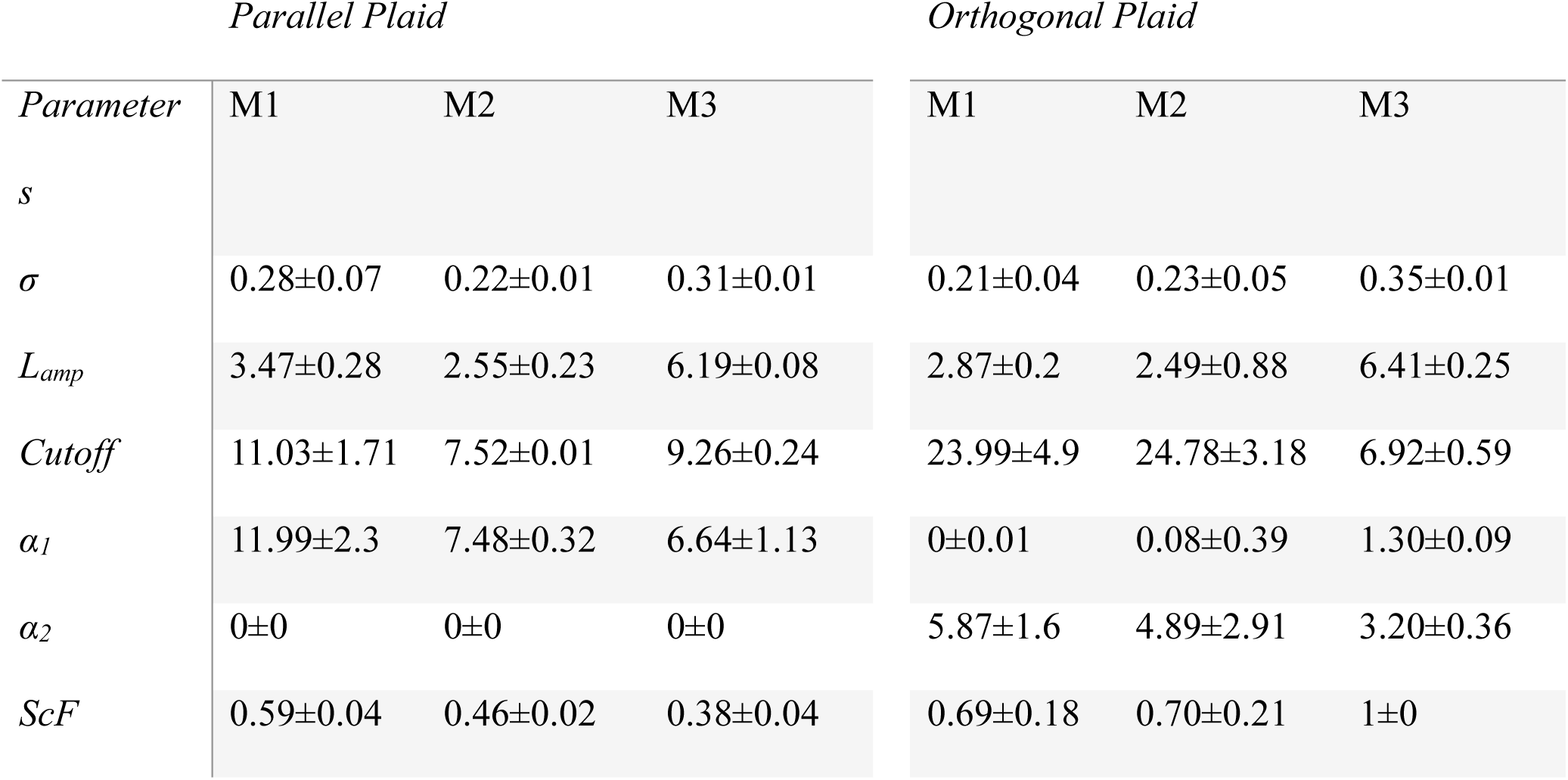
Showing median ± bootstrapped SEM values for parameters obtained from the optimal-tuned normalization model for LFP and ECoG data.

**Table 5:** Showing median ± bootstrapped SEM values for parameters obtained from the optimal-tuned normalization model fitted for spiking data.

|  | *Parallel Plaid* | *Orthogonal Plaid* |
| --- | --- | --- |
| *Parameters* | M1 and M2 | M1 and M2 |
| $\sigma$ | 0.48±0.14 | 0.40±0.15 |
| $L_{amp}$ | 5.98±0.41 | 6.45±2.13 |
| *Cutoff* | 7.83±0.58 | 15.97±2.27 |
| $\alpha_1$ | 8.15±2.71 | 0.03±0.37 |
| $\alpha_2$ | 0±0.99 | 2.43±1.88 |
| *ScF* | 0.32±0.26 | 1±0 |

**Table 6:**
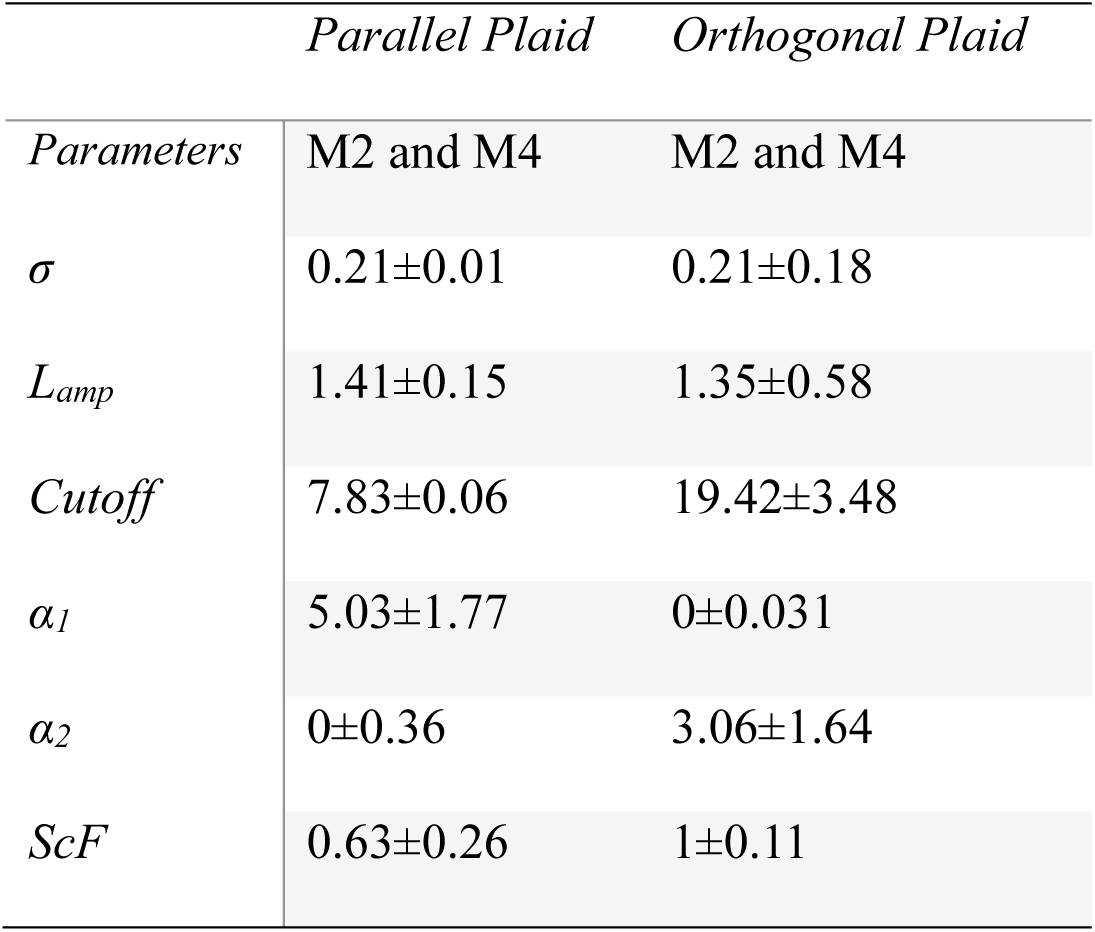
Showing median ± bootstrapped SEM values for parameters obtained from the optimal-tuned normalization model fitted for EEG data.

|  | *Parallel Plaid* | *Orthogonal Plaid* |
| --- | --- | --- |
| *Parameters* | M2 and M4 | M2 and M4 |
| $\sigma$ | 0.21±0.01 | 0.21±0.18 |
| $L_{amp}$ | 1.41±0.15 | 1.35±0.58 |
| *Cutoff* | 7.83±0.06 | 19.42±3.48 |
| $\alpha_1$ | 5.03±1.77 | 0±0.031 |
| $\alpha_2$ | 0±0.36 | 3.06±1.64 |
| *ScF* | 0.63±0.26 | 1±0.11 |

The fit quality obtained from the original-tuned model was slightly better than that of the optimal-tuned normalization model across electrodes and monkeys for both parallel and orthogonal plaids. (Figure 7A Wilcoxon signed rank test, parallel: p = 2.463e-14; orthogonal: p = 5.215e-13). When differences in model parameters were compared, the corrected Akaike Information Criteria values (AIC_C_) remained significantly higher for the optimal-tuned normalization model (Wilcoxon signed rank test, parallel: p = 5.570e-10; orthogonal: p = 0.0018; Figure 7B).

**Figure 7.**
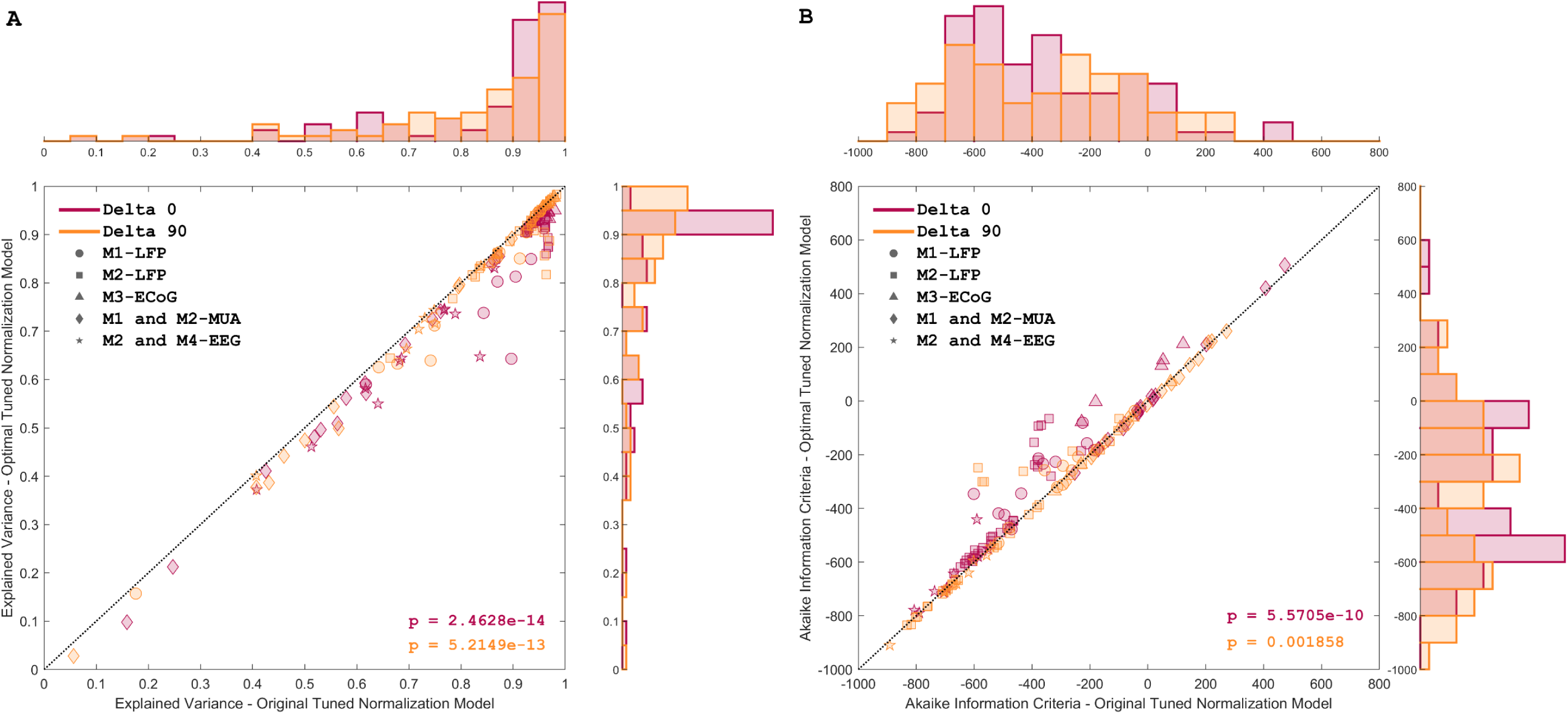
Comparison of performance for original-tuned and optimal-tuned normalization model. Scatter plots with histograms for (A) explained variance and (B) Akaike information criteria calculated for original-tuned and optimal-tuned normalization models. The subplots at the top and right represent the histograms of counts for each model. Each data point in the scatter plot represents one electrode. Colors denote parallel (deep magenta) and orthogonal (gold) plaid conditions. Different markers are used to indicate the values corresponding to spiking, LFP, ECoG, and EEG fits obtained across the four monkeys. P-values obtained from the two-tailed Wilcoxon signed rank test are denoted in the bottom left.

While the optimal-tuned normalization model achieved the desired suppression profiles with very few parameters, we could not explain the asymmetry in the profile near the target frequency (compare the magenta traces in Figure 4A versus 6A), which could also have explained the slightly poorer performance of the optimal-tuned normalization model compared to the original model. However, we found that such asymmetry in the suppression could be achieved by passing the inhibitory drive from non-linearity of different order other than squaring (Figure S3), as elaborated in the <u>Discussion</u>.

### Difference-Intermodulation terms are also low-passed

We also studied the sum and difference intermodulation (IM) terms in LFP and ECoG as a function of the contrast and temporal frequency of the mask grating (Figure S4 and S5). IM terms were predominantly present for parallel plaids and became stronger as the mask and target contrast were increased. Similar results were observed in EEG data as well (not shown). The terms were barely perceptible for orthogonal (Figure S4 panel D-F) and 45° separated plaids (Figure S5) except when the temporal frequency of the mask was a harmonic of the temporal frequency of the target. Again, this indicates that the temporal frequencies pass through the non-linear stage together when the orientation of the two gratings matches. The difference-intermodulation terms (*f_1_-f_2_*) had the minimum amplitude response when the temporal frequencies of the target and mask grating were closest, and the response increased as the two moved away (Figure S4, panel B). As before, this could be attributed to the low-passed normalization drive, which becomes strongest when |ω_*T*_ − ω_*M*_| is small, thereby leading to a small amplitude change. The results for sum-intermodulation components are discussed in the supplementary.

### Additional evidence for low-pass filtering

#### The preferred temporal frequency of a grating depends on stimulus size

If the normalization signal has more energy from lower frequencies, we hypothesized that the temporal frequency tuning of counterphase gratings should be a function of stimulus size. When a larger stimulus is presented, the feedforward input remains the same as when a small stimulus is presented, but since more surround is activated, it leads to a more extensive interneuronal network activation^31^. A recent study has also shown reduced synaptic excitatory input for a larger stimulus compared to a smaller stimulus^32^. Thus, the normalization signal would be more robust for larger stimuli, and therefore, the lower temporal frequencies would be suppressed more for larger stimuli compared to small stimuli. This, in turn, will lead to a higher preferred temporal frequency for larger stimuli. To test this, we presented counterphase gratings of two sizes (full-field and 1.5°) at various temporal frequencies (Figure 8). We observed that for a maximum contrast small-sized stimulus, a clear peak in the tuning curve was indiscernible, with 4Hz-16Hz having approximately a similar amplitude change. However, a distinguishable peak at 16Hz was evident for the large stimulus. Consistent with the hypothesis, the preferred TF shifted towards higher frequencies when stimulus size was increased across all stimulus contrasts (Figure 8D; right-tailed Wilcoxon signed rank test, p = 1.8199e-14).

**Figure 8.**
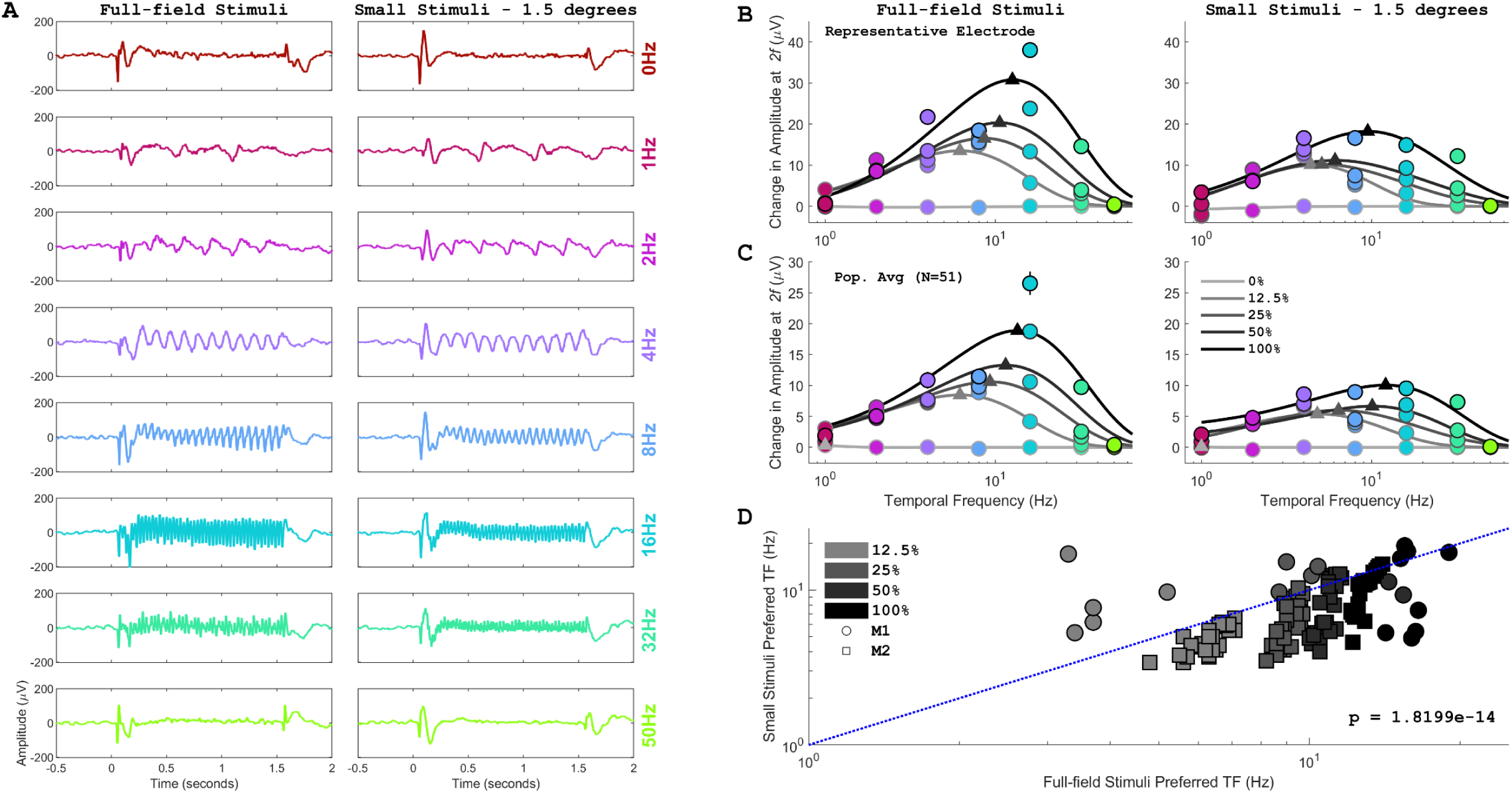
SSVEP tuning profiles of counterphase gratings in the LFP signal as a function of stimulus size. (A) Evoked responses of a representative electrode in response to a full-field and a small (1.5°) counterphasing grating stimulus presented at varying temporal frequencies (0,1,2,4,8,16,32, and 50Hz) and 100% contrast. (B) SSVEP tuning plots show the change in amplitude at twice the temporal frequency of the stimulus as a function of contrast (light to dark shaded markers) plotted for the representative electrode for the two stimulus sizes. The responses for each contrast were fitted using the difference of exponentials function (equation 2). Triangles represent the preferred TF. (C) Same as (B) but for LFP responses averaged across sessions and electrodes for M1 and M2. ‘N’ indicates the total number of electrodes across the two monkeys. (D) Scatter plots of preferred TFs estimated for full-field and small stimulus for each electrode of M1 and M2. Each data point represents an electrode with fit quality >= 0.6 and a preferred TF > 1Hz for a particular contrast condition (grey colour levels).

## DISCUSSION

By using counterphasing plaid stimuli, we demonstrated that the masking interactions in V1 neurons are a function of both temporal frequency and relative orientations of component gratings, which holds irrespective of the stimulus size and extends to multiple recording scales, including ECoG and EEG. The band-pass suppression profile obtained for parallel plaids and the low-pass suppression profile obtained for the orthogonal plaids can be explained simply by adding a low-pass filter to the normalization pool, which receives input from orientation-selective channels. We also show that the rightward shift of the preferred temporal frequency of a counterphase grating when stimulus size increases and the amplitude of difference-intermodulation terms as a function of the temporal frequency of the mask can also be attributed to low-pass filtering of the inhibitory drive.

### Comparison with previous studies

Divisive normalization has significantly explained pattern masking in the temporal domain. While some studies^7–9^ have assumed the model to be memoryless, many previous studies have used low-pass filters to explain the dynamics of normalization^6,16–18^. However, most studies have used the filter to explain the temporal dynamics of the response. We have used the low-pass filter to instead explain the shape of the suppression function, not the temporal dynamics of the response. Specifically, the response and suppression profiles in equation 4 and 5 are not functions of time. Although the computation of S is done by low-pass filtering signals that are functions of time, we averaged the energy over time (which can be thought of as another ‘hidden’ low-pass filter with a large time-constant in our formulation). Since the characteristics of the low-pass filter were not used to explain the temporal variation of the response, this is an independent approach to study the characteristics of the filtering action.

Reassuringly, this alternate approach also yielded compatible results. We obtained cutoff of the low-pass filter between ∼7-11 Hz for parallel plaids and ∼15-30 Hz for non-parallel plaids (except for M3, where the cutoff was ∼6 Hz). These correspond to an integration time (1/(2π *f*)) of the normalization signal to be about ∼14-23ms and ∼5-10ms for parallel and orthogonal plaids, respectively. Note that for the orthogonal condition, the filtering action produces a “low-pass” effect, with shallower slopes corresponding to high filter cutoffs. However, the same can be achieved by having a large value of the semi-saturation constant, leading to a shallower overall slope. The parallel condition puts stronger constraints on the low-pass filter, for which the time constants (14-23 ms) were in agreement with previous reports that have reported time constants of ∼25-30ms in humans^6^ and monkey V1^19^ and between ∼5-25ms in cat’s retinal ganglion cells^33^, albeit the mechanisms that give rise to these time constants are very different (as discussed below).

However, it is unlikely that a single low-pass filter with a cutoff of ∼10 Hz is sufficient to explain the temporal dynamics of normalization. Note that a counterphase grating, even at low frequencies, produces a response only at the counterphasing frequency and its harmonics. If the normalization signal which comes in the denominator of equation 4 is allowed to oscillate, the resulting response function will have many other frequencies since division by a sinusoid generates a non-sinusoidal signal. Since this is not observed in real data, the normalization signal must be low-pass filtered with a sufficiently long time constant (in our formulation, the averaging operation in equation 5 was this ‘hidden’ filter). For example, Zhou and colleagues^17^ found a time constant of ∼100 ms when they fitted the temporal dynamics of the response. It is likely that multiple filters are operating at different levels, as suggested previously^33^, which could explain stimulus interactions as well as the temporal evolution of the response.

Our results are more comparable with those of Tsai and colleagues^6^, who also used a low-pass filter to explain the changes in responses at target, mask and intermodulation frequencies with varying contrasts. However, contrast response functions were only obtained for a single target and mask frequency, as in other studies as well^6,8,28^, whereas we obtained contrast responses for the target against multiple temporal frequencies and orientations of mask grating. Our time constants match those reported by Tsai et al., even though the stimulus used was a random noise pattern, unlike the counterphase plaids we used.

For orthogonal plaids, we found a low-frequency tuned suppression at lower contrasts that changed to non-specific suppression at high contrast. This can happen because the relative contribution of the suppression term in the normalization signal reduces when the target contrast is high (see equation 5). Note that we kept the semi-saturation constant fixed; in a previous report, it was varied as the function of mask contrast^6^. So, the change in the response profile at different contrast levels could also be explained by varying the semi-saturation value with contrast. For example, if the semi-saturation constant increases with target contrast, it could also make the suppression profile non-specific at high target contrasts.

According to the winner-take-all computation, the stimulus with higher contrast should be favoured for combinations where the target or mask stimuli have dissimilar contrasts^4^. This would also mean that IM components should be negligible for such cases, as shown by Tsai and colleagues^6^. However, in our results, we did not observe any enhancement in IM components when target and mask contrasts were matched (Figure S4; matched target and contrast conditions correspond to light green in the first column of S4B (both at 0%), dark green in second (both at 6.25%) and so on, which did not seem out of place compared to the remaining non-matched conditions). These observations were in contrast to those observed by Tsai et al., as they observed sum-IM components decreasing with increasing mask contrast beyond the maximum target contrast, and they did not obtain strong difference-IM components^6^. However, they used modulation frequencies for the two stimuli very close to each other, with a difference of ∼2Hz (|ω_*T*_ − ω_*M*_|), comparable to the data points at mask frequencies of 13 or 17 Hz in Figure S4B-S4C. We also found the difference IM responses to be small for that case, since the IM components were able to pass through the low-pass filter and contribute in the suppressive drive. By using a much wider range of mask frequencies compared to previous studies, a more complete IM interaction profile could be observed.

We observed that IM terms were absent from the responses for orthogonal and 45° separated plaids and were found only for parallel plaids, hinting that the orientation bandwidth for the filters is less than 45 degrees. Results from a previous EEG study^8^ also align with this observation, where the presence of IM terms falls to negligible at 45° separation. This is not surprising since the orientation channel bandwidth in V1 is reported to be around ∼20-30° in neural^34^ and psychophysical responses^35^. Therefore, 45° separated plaids are likely to be processed by separate filters and, hence, are supposed to be more similar to the orthogonal case than the parallel one, as observed in our data.

### Underlying architecture of slow-varying normalization

The stimulus sizes we used (1-1.5° and full-field) activate two kinds of suppression – overlay and surround suppression^36^. Suppression is generally delayed by ∼10-25ms with respect to the onset of excitatory response^37–39^, although the delay is dependent on the strength of suppression^38^. Overlay suppression is relatively faster by ∼10ms than surround suppression^39^ but is also broadly tuned^36^. Contrarily, near and far surround suppression are orientation-tuned and arise from lateral and feedback connections, although near surround suppression is much more sharply tuned^38,40^. This suppression of responses because of masking is mediated by GABA_A,_ irrespective of the relative orientation difference between the plaid components^41^. GABAergic neurons, which include somatostatin-expressing (SST) and parvalbumin-expressing fast-spiking (PV) interneurons, are thought to play distinct roles in normalization^42^, input summation and output normalization. SST neurons have slower dynamics but are selective, while PV neurons provide faster-untuned feedforward suppression^43,44^. Since orthogonal suppression becomes non-specific at higher contrasts and has relatively faster time constants, it may have a higher contribution from PV interneurons. On the other hand, SST neurons might play a more prominent role in the suppression caused by parallel plaid stimuli. The onset latency of SST neurons is about ∼20-25ms delayed compared to that of pyramidal and PV cells^43^; this is equivalent to the time constants we obtained from the model for parallel plaids.

A recent study^45^ showed that the frequency fall-off in gamma oscillations arising from such excitation-inhibition interactions could also be attributed to delayed surround input. The study used an inhibitory stabilized network (ISN) model, where delayed inhibition was modelled as Ornstein-Uhlenbeck (OU) noise, which can be thought of as Gaussian white noise passed through a low-pass filter. Similar ISN models have also been used to explain masking dynamics^46,47^. Thus, there are multiple ways to incorporate slow dynamics in the inhibitory circuit, and more work is warranted to explain the mechanisms that lead to temporal frequency masking interactions described here.

### Model peculiarities and limitations

The optimal-tuned normalization model successfully explained the overall profile of the suppression but failed to capture the asymmetry observed in the suppression profile for the parallel plaids. One way to achieve asymmetry using the current form of the model would be to change the exponent when the inhibitory drive passes through the orientation filter. With even exponents (Figure S3 B and D), the band-pass profile obtained is symmetric as |ω_*T*_ − ω_*M*_| is the same for both sides of the target frequency. However, with odd exponents, an asymmetric low-pass profile emerges where the temporal frequencies lower than that of the target suppress more. This happens because the Fourier domain components are no longer symmetric, and energy found for lower mask frequencies comes closer to the cutoff of the low-pass and can, therefore, pass through it (Figure S3, panels A and C, third column, lavender peaks). Hence, contributions from the discrete populations of V1 neurons that follow odd and even-order non-linearities can give rise to such asymmetry in the response.

Interestingly, the low-pass suppression profile that we observed for orthogonal plaids can also be obtained by using odd exponents in the model (Figure S3 panel A and C). However, this would suggest that, like parallel plaids, these stimuli also pass through the same filters before any non-linearity. Some studies^48,49^ have favoured this hypothesis for target and mask stimuli at different orientations. However, unlike our study, they used stationary stimuli at different spatial frequencies, but we matched the spatial frequencies of the component gratings of the counterphasing plaid. Studies^8,28^ in the past, which have used orthogonally oriented temporally modulated stimuli, favoured passing signals through non-linearity and then summing them up in a way similar to that obtained by our model. For orthogonal plaids, the suppression profile for M3, which was computed using ECoG signals, and the EEG profile computed for M2 and M4 were different from the LFP and spiking profiles obtained for M1 and M2 (Figure 6). It reflected contributions from both hypotheses but had a higher contribution from the addition of non-linear signals (α_2_ > α_1_, Table 4 and Table 6). The difference could simply be due to the larger spatial summation of ECoG and EEG signals compared to LFP^50^. In summary, we note that adding more filters, making the semi-saturation constant a function of contrast, or changing the exponent can all lead to improvements in the model to capture additional features (e.g., temporal dynamics of the response or asymmetry in suppression profile).

### Ramifications for cognitive and BCI studies

Our findings underscore the need to carefully account for changes in SSVEPs that are purely driven by sensory interactions while using them to study various cognitive tasks, such as visual attention, working memory, and binocular rivalry^22,25,51,52^. This emphasizes the need to carefully consider stimulus parameters while designing studies and choose parameters that have minimal sensory interactions to delineate frequency and feature-driven effects. However, since EEG signals come from multiple visual areas, whether higher visual areas also show similar dynamics needs to be examined. Additionally, the receptive fields of higher visual areas tend to be much larger than V1; therefore, the effects of non-overlapping but concurrently presented stimuli can be systematically investigated, which was not feasible for the current recordings. Our findings could also help enhance the precision and reliability of BCIs, which often rely on carefully measuring SSVEP responses. It has also been observed that the performance of BCI is often contingent upon features such as colour and spatial proximity of stimulus^53,54^, so adapting our current paradigm to study these features may yield valuable insights into the underlying neural circuitry and how to optimize such designs.

## METHODS

All the animal surgeries and experiments were reviewed and approved by the Institutional Animal Ethics Committee of the Indian Institute of Science and were conducted following the guidelines approved by the Committee for the Purpose of Control and Supervision of Experiments on Animals (CPCSEA). Four adult bonnet macaques (*Macaca radiata*; three females and one male) participated in the experiments (M1 – 3.2 kg, 13 years; M2 – 5.8 kg, 15 years; M3 – 3 kg, 17 years; M4 – 6.7 kg, 18 years).

### Animal Preparation and Training

Before training, each monkey was surgically implanted with a titanium headpost over the frontal region of the skull under general anaesthesia. Following a recovery period, monkeys were trained on passive fixation tasks. After the monkeys learned the task, a second surgery was performed to implant a microelectrode grid (Utah Array, Blackrock Neurotech). Grids consisted of platinum microelectrodes of 1mm in length, with a tip diameter of 3-5µm and an interelectrode distance of 400µm; the grids were centered ∼10-15mm rostral from the occipital ridge and ∼10-15mm lateral from the midline, with slightly varying locations across the monkeys. M1 was implanted with a 10×10 microelectrode array grid with 96 active electrodes, and M2 was implanted with a 6×8 microelectrode grid with 48 active electrodes in the right primary visual cortex. In M2, a second grid of similar specifications was implanted in the V4 region, but this study did not analyze those signals. M3 was implanted with a custom hybrid array that consisted of a 9×9 microelectrode grid with 81 active microelectrodes and a 3×3 electrocorticogram (ECoG) grid (Ad-Tech Medical Instrument Corporation). ECoG contacts were 2.3 mm wide platinum discs with a 10mm center-to-center interelectrode distance. The current study analyzed only ECoG electrodes, as the microelectrode grid had stopped working before recording the experiments described in this study. M1 and M3 were also used in our previous findings on SSVEP local interactions, as monkey two and monkey one, respectively^10^. Reference wires were inserted under the dura at the edge of the craniotomy or secured to titanium screws or metal straps, which were used to secure the bone of the craniotomy. Referencing was performed using a 96-channel Cereplex E headstage. The receptive field of the recorded neurons was located in the lower left quadrant for M1 and M2 and the lower right quadrant for M3, about the fixation point (at the eccentricity of ∼2.7°-4.4° in M1, ∼2.6°-3.3° in M2, and ∼1.9°-7.2° in M3, Figure S1). Neural data was collected once monkeys recovered from the grid implantation surgery and performed the task. In addition, 18-channel electroencephalogram (EEG) data were collected from M2 and M4. We used Ag-AgCl cup electrodes for EEG recordings, which were secured on the monkey’s scalp using a conductive paste (Ten20 conductive electrode paste, Weaver and Company). Before putting the electrodes, a mild abrasive gel was used to lower the impedance (NuPrep Skin Prep Gel, Weaver and Company).

### Experimental Setup and Data Acquisition

Monkeys sat ∼50cm away from the monitor in a primate chair, their heads restrained by a headpost. The monitor (BenQ XL2411, LCD, 1280×720 resolution with a refresh rate of 100Hz) was gamma-corrected using Colorchecker Display Plus (Calibrite; formerly known as i1 Display Pro), with the gamma value set to unity and mean luminance set at 60 cd/m2. Monkey’s eye position data (horizontal and vertical coordinates/positions and pupil diameters) were tracked and recorded using the ETL-200 Primate tracking system (ISCAN, sampled at 200Hz). The display setup, eye tracker, and the animal were housed in a dark Faraday enclosure lined with a copper sheet to minimize external electrical interference. Stimulus generation, pseudorandom stimulus presentation, and eye signal monitoring were done using custom software (Lablib) running on macOS. Neural signals, local field potentials (LFP), and multiunit activity (MUA), were recorded using a 128-channel Cerebus neural signal processor (Blackrock Neurotech). Raw data was band-pass filtered between 0.3Hz (analog Butterworth filter, first order) and 500Hz (digital Butterworth filter, fourth order) and was sampled at 2000Hz, digitized at a resolution of 16 bits to collect LFP, ECoG, and EEG data. To obtain multiunit activity, neural signals were sampled at 30KHz and filtered between 250Hz (digital Butterworth filter, fourth order) and 7500Hz (analog Butterworth filter, third order), followed by amplitude thresholding the signal at ∼5 standard deviations below the mean (except for one session for M2, where it was kept at 4). The MUA was not sorted further, and no further offline filtering was done on the data. For EEG, the ground electrode was placed right lateral to the headpost, and the reference electrode was placed behind the headpost.

### Experimental Design and Visual Stimuli

Each experiment started by presenting a fixation dot (radius 0.1°) at the center of the display monitor. Monkeys were trained to fixate on this dot while holding their gaze within a fixation window (1.5° for M1, M2, and M4, and 2° for M3) for the entire trial duration to get a juice reward at the end. Monkeys were given ∼1000-2000ms to start fixating; if they failed to fixate within this stipulated time or broke fixation during the trial duration, the trial was aborted. While monkeys fixated, a series of achromatic counterphasing gratings or plaid stimuli (generated by overlapping two counterphasing gratings) were presented. We ran three experiments with different stimulus presentations.

In Experiment 1 (Figure 2), performed by M1 and M2, full-field counterphasing plaid stimuli were pseudorandomly presented. The trial began with a fixation period (1000ms for M1 and 1500ms for M2) followed by two stimulus presentations of 1500ms each, with an inter-stimulus interval of 1500ms. The inter-trial interval was kept at 1500ms (for M1) or 1000ms (for M2). The first grating (the target grating) had an orientation of either 0° or 90° in a session with a fixed temporal frequency of 15Hz. The second grating (the mask grating) was presented at two different orientations (0° and 90°) at varying temporal frequencies (1-29Hz in steps of 1Hz). The contrast of the target grating was either kept at 0 or 25%, and the mask grating had a fixed contrast of 25%, allowing us to record responses to grating and plaid stimuli in the same session. The spatial frequency of both gratings was fixed at four cycles/degree (cpd). Each session had 116 stimulus conditions (2 target contrasts × 2 mask orientations × 29 mask temporal frequency), with at least 15 repeats for each stimulus. We collected data from one session for M1 and two sessions for M2.

Since full-field stimuli highly suppress MUA response; in Experiment 2 (Figure 2-7), a smaller stimulus (radii -1° for M1 and 1.5° for M2) spanning the aggregate receptive field of the microelectrodes was presented. Target and mask gratings were presented at four contrasts (0 to 25% on a log scale) at a spatial frequency of 4 cpd. Superimposing the two gratings doubled the contrast of the resulting plaid stimulus. Multiple contrasts allowed us to estimate the parameters for the normalization model (<u>explained below</u>) as the complete profile of masking responses could be obtained. The temporal frequency of the target grating was fixed at 15Hz, while the mask grating in a trial could take any value between 1-29Hz (in steps of 2Hz). Orientation of target or mask grating could either be at 0° (parallel plaids) or 90°(orthogonal plaids) for M1 or could take any value between 0° and 135° (in steps of 45°) for M2, such that the difference between the orientation of the two gratings is either 0° or 90° for a single session. A total of 240 unique stimulus conditions (4 target contrasts × 4 mask contrasts × 15 mask temporal frequency) were repeated 15 times in one session. Since we had almost doubled the stimulus conditions compared to the first experiment, we reduced the stimulus duration to keep the time taken to finish a session the same, as it was already between ∼2.5-3 hours. Three stimuli of 800ms each were presented in a trial after an initial fixation period of 1000ms with an inter-stimulus interval of 700ms, followed by an inter-trial interval of 1000ms. We collected data from 2 sessions from M1 (one session each for parallel and orthogonal plaid) and across 8 sessions (4 sessions each for parallel and orthogonal plaids) for M2. To optimize the stimulus parameters for recording ECoG activity from M3 and EEG activity from M2 and M4, we ran the same experiment with a full-field stimulus instead, with a target and mask orientation of 0° or 90°. From M3, we collected data from 3 sessions for orthogonal plaids and 2 sessions for parallel plaids. For EEG data, one session for each parallel and orthogonal plaids was recorded from both M2 and M4. In M2 and M3, we also ran additional sessions, where the orientation difference between the target and mask gratings was fixed at 45° (4 sessions for M2 and 2 sessions for M3, Figure S2).

Experiment 3 (Figure 8) involved presentations of counterphase gratings of full-field and 1.5° in size across five contrast levels (0 to 100% on a log scale) and 8 temporal frequencies (0, 1, 2, 4, 8, 16, 32, and 50Hz). The orientation and spatial frequency of gratings were fixed at 90° and 4cpd. Each trial started with a fixation period of 1000 or 1500ms, followed by two or three stimulus presentations. Stimulus was presented for 800ms for M1 and 1500ms for M2 with an interstimulus interval of 700 and 1500ms, respectively. Eighty stimuli (2 sizes × 5 contrasts × 8 temporal frequency) were each repeated at least 12 times. Recordings were done from one session for M1 and two sessions for M2.

### Receptive Field Mapping and Electrode Selection

A receptive field mapping protocol, which involves the flashing of small grating stimuli (0.3°-0.4°) on equally spaced locations within a rectangular grid (11×11 for M1, 14×17 for M2, and 19×25 for M3), was run across multiple days to estimate receptive field locations of the electrodes. While the monkey fixated on the central fixation spot, a series of stimuli, each for 200ms (with an interstimulus period of 0ms), was presented at full contrast. The stimulus had a spatial frequency of 4cpd (for M1 and M3) or 2cpd (M2) and could take any of the four orientations (0°, 45°, 90°, or 135°). The evoked LFP responses were used to estimate receptive field centers and sizes by fitting a 2D Gaussian. Any electrode with a stable receptive field center across 50% of the sessions and a receptive field size within 0.1° to 0.5° was selected for LFP analysis. Impedances for all the electrodes across all sessions were under 2500kΩ. This yielded 21 and 32 microelectrodes for M1 and M2, and 5 ECoG electrodes for M3. In M1, for experiment 2, the recording quality was relatively poor, so only electrodes with strong evoked responses in that session (N = 10-11 electrodes) were used. For MUA analysis, we first selected electrodes that accumulated more than a certain number of spikes in one session (>= 4000) and had a signal-to-noise ratio (SNR) and change in firing rate more than a predetermined cutoff (SNR >= 1.2, firing rate during stimulus period (250-750ms) >= 1 spikes/second and had a sharp transient increase of activity during first 150ms of stimulus presentation were used (greater than 1.5 times the activity during baseline period). We obtained 79 electrodes for the parallel plaids session and 56 for the orthogonal plaids session, pooled across both monkeys (M1 and M2) across all the sessions. All these units were visually active, but only a few had high sustained firing rates. We further subselected units which had more stringent cutoffs (# of spikes in a session >=10000, SNR >=1.8, Firing rate during stimulus period >= 6spikes/sec, and had a sharp increase in transient). This way, we obtained 18 and 15 electrodes for parallel and orthogonal plaids sessions, pooled across both monkeys. For EEG analysis, after removing electrodes with high impedance, electrodes with a more than 0.5μV change in amplitude during stimulus period from baseline period (equation 1) at 30Hz were used for analysis. This yielded a total of 11 and 7 electrodes for parallel and orthogonal plaids sessions from M2 and M4, located at and around the occipital region.

### Data Analysis

All the data were analyzed using custom-written codes in MATLAB (MathWorks, RRID: SCR_001622). For the analysis window, the stimulus period was defined as 250 to 750ms or 250 to 1250ms post-stimulus onset, depending on the stimulus presentation duration. The baseline was defined as the corresponding period before the stimulus onset (-500 to 0ms or - 1000 to 0ms). Consequently, this results in a frequency resolution of 2Hz or 1Hz. Trials with excessive noise (due to any movement or electrical artefact) during these periods were discarded if the amplitude of the waveform exceeded a threshold of six times the standard deviation from the mean waveform. Thereby rejecting <2% of trials for each monkey for all experiments. The remaining waveforms were then used for further analysis. The counterphasing stimulus evokes a more vigorous SSVEP response at the second harmonic, and the amplitude difference was calculated at twice the temporal frequency (*f*) of the grating (for experiments 1 and 2 – at 2*f of* target grating, i.e. 30Hz, experiment 3 – at 4, 8, 16, 32, 64 and 100Hz) between stimulus and baseline period (equation 1). For experiment 2, the change in amplitude was also calculated at intermodulation terms (*f_1_-f_2_* and *f_1_+f_2_*). For LFP, ECoG, and EEG analysis, the baseline across all stimulus repeats was averaged, and a common baseline was used. Amplitude spectra were calculated for the stimulus and baseline period using the discrete Fourier transform for individual trials. To obtain firing rate spectra, we generated peristimulus time histograms by binning spikes across each trial in 10-ms bins and then averaging across trials for each condition, followed by their Fourier transform.

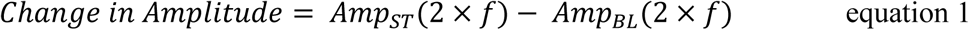

SSVEP tuning functions (Figure 8) were fitted using the difference of an exponential function^55,56^ (equation 2) using an ordinary least squares fit. The peak of the fitted function was considered as the preferred temporal frequency. Electrodes were classified as low-pass if the preferred temporal frequency of an electrode was <=1 Hz and otherwise called band-pass. The statistics of preferred temporal frequency are calculated further only on band-pass with a fit quality of >=0.6.

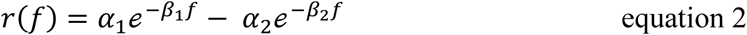

*r*(*f*) is the change in amplitude at twice the temporal frequency of the grating. Coefficients α1, β1, α2, and β2 were constrained to be non-negative. The fit quality (*qval*) was calculated as the fraction of the total variance in the data explained by the fits:

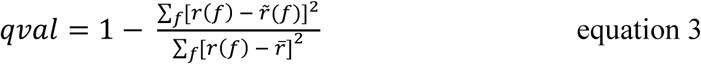

where *r̃*(*f*) is the fitted value corresponding to *r*(*f*) and *r̅* is the mean change in amplitude across all temporal frequencies. A qval close to 1 corresponds to decent fits, whereas negative qval values close to 0 indicate poor fits.

### Normalization model

A tuned divisive normalization model, which has been previously used to explain SSVEP interactions^9^, was adapted to explain the suppression profile obtained for the amplitude of the target grating as a function of the temporal frequency of the mask grating (equation 4, Figure 1 panel A) for experiment 2. The following equation describes the model:

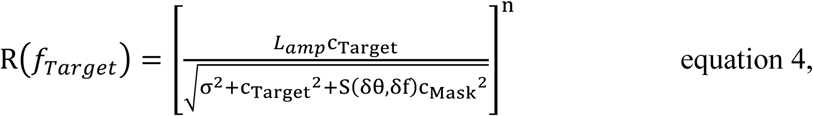

where R(*f_Target_*) is the resultant mean amplitude response at twice the temporal frequency of the target (30Hz). L_amp_ corresponds to the response of an individual grating at the unit contrast, σ, and n represents the semi-saturation constant and non-linearity in the gain of the response. S(δθ,δf) captures the suppression driven by the mask grating with a particular orientation and temporal frequency. First, we kept the following sixteen as non-negative free parameters: L_amp_, constrained between 1 and 15; σ, constrained between 0.1 and 4; and S (14 mask temporal frequencies), constrained <35. Exponent ‘n’ was fixed at a value of 2. This version of the model is referred to as the original-tuned normalization model in the rest of the text.

A second, optimal-tuned normalization model was also fitted, which accounts for a functional form for ‘S’. In this approach, we assumed that the neural responses towards the target and mask grating are sinusoidal. Similar to the Foley-style model^27^ (equation 8 - Model 3), we presumed that depending upon the orientation difference between the target and the mask grating, the component sinusoid contribution would be summed before or after the exponentiation. We also argue that the normalization signal, coming as the inhibitory drive, will vary slowly over time, thereby integrating responses from the pool within a temporal window. This can be achieved by low-pass filtering (LPF) of the signal, similar to the previous studies^6,17^. In this model, ‘S’ was assumed to be the mean energy of the inhibitory gain pool and was denoted as:

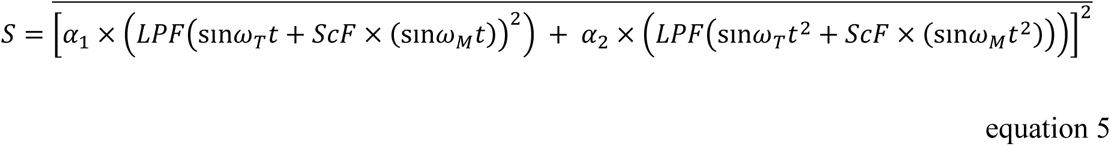

where ꞷ_T_ and ω_M_ represent the angular frequency of the target and mask grating. α_1_ and α_2_ represent the weights of the low-pass drive of the two orientation-selective filters. The bar denotes average over time (so S is not a function of time). The scaling factor (*ScF*) factors in the contribution from the mask grating as the signals from the target and mask might be weighted to allow an optimal combination. It primarily controls the contribution from lower mask frequencies. We used a fourth-order Butterworth low-pass filter and left the filter cutoff as a free parameter. But before passing the signal through the low-pass filter, we corrected the signal to remove the DC component. We plugged the value of S obtained from this equation back into equation 4 to estimate the final response. In this model, we had fixed the exponent at two and had six non-negative free parameters: L_amp_, constrained between 1 and 15; σ, constrained between 0.1 and 4; cutoff frequency of the filter, constrained between 1 and 40; α_1_ and α_2,_ constrained >0_;_ and *ScF*, constrained between 0.1 and 1.

The curve fitting was done using the fmincon function in MATLAB to minimize the sum of the squared estimate of the errors:

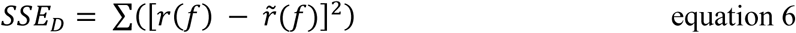

where *r*(*f*) is the observed response and *r̃*(*f*) is the fitted value. The quality of the fits provided by the model was assessed by computing the explained variance (equation 3).

We used corrected Akaike information Criteria^57,58^ (AIC_C_) to compare the model performance as the two models varied in their complexity owing to different numbers of free parameters.

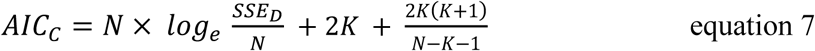

where N refers to the number of observations, SSE_D_ is the sum of the squared estimate of errors (equation 6), and K represents the number of free parameters in the model. We also added SSE_D_ as an additional free parameter^58^ for the calculation, increasing the tally of free parameters to 17 and 7, respectively, for the two models. The AIC_C_ values are comparative in nature and do not have any interpretation on their own.

### Quantification and statistical analysis

We performed a two-tailed Wilcoxon signed rank test to compare the model performance across two versions of the normalization model, with the null hypothesis that samples come from distributions with equal medians. We used the right-tailed Wilcoxon signed rank test to compare whether the preferred temporal frequency changes with stimulus size, with the null hypothesis that larger stimuli prefer faster temporal frequencies than small ones.

## Author Contributions

Conceptualization, D.G., S.R.; methodology, D.G., S.R.; investigation, D.G.; formal analysis, D.G., S.R.; writing – review and editing, D.G., S.R.; funding acquisition S.R.; supervision S.R.

## Funding Disclosure and Acknowledgements

This work was supported by Wellcome Trust/DBT India Alliance (Senior Fellowship IA/S/18/2/504003) and DBT-IISc Partnership Programme to S.R. and Senior Research Fellowship from Council of Scientific and Industrial Research to D.G.

## Data and Code Availability

Data and codes are available on https://osf.io/jzb5u/ and https://github.com/Divya-Gulati/Temporal-Frequency-Masking-, respectively.

## SUPPLEMENTAL INFORMATION

### Figure Legends

**Figure S1.**
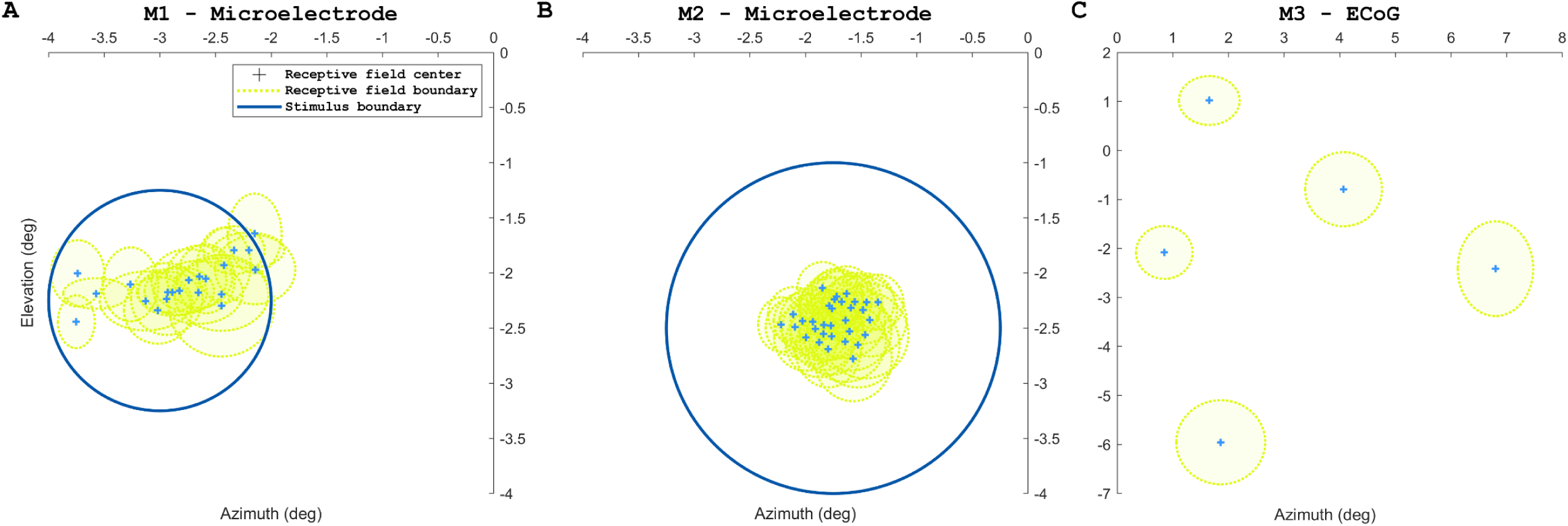
Location of receptive fields and stimulus. Locations of receptive field centers with their respective sizes of the microelectrodes implanted in V1 of (A) M1, (B) M2. Only electrodes with stable receptive fields across days are plotted. Refer to <u>receptive field mapping and electrode selection</u> in the <u>methods</u> section for more details. The blue circle marks the size and location of the stimulus presented in <u>experiment 2</u>. (C) Locations of receptive field centers with their sizes for ECoG grid implanted in M3.

**Figure S2.**
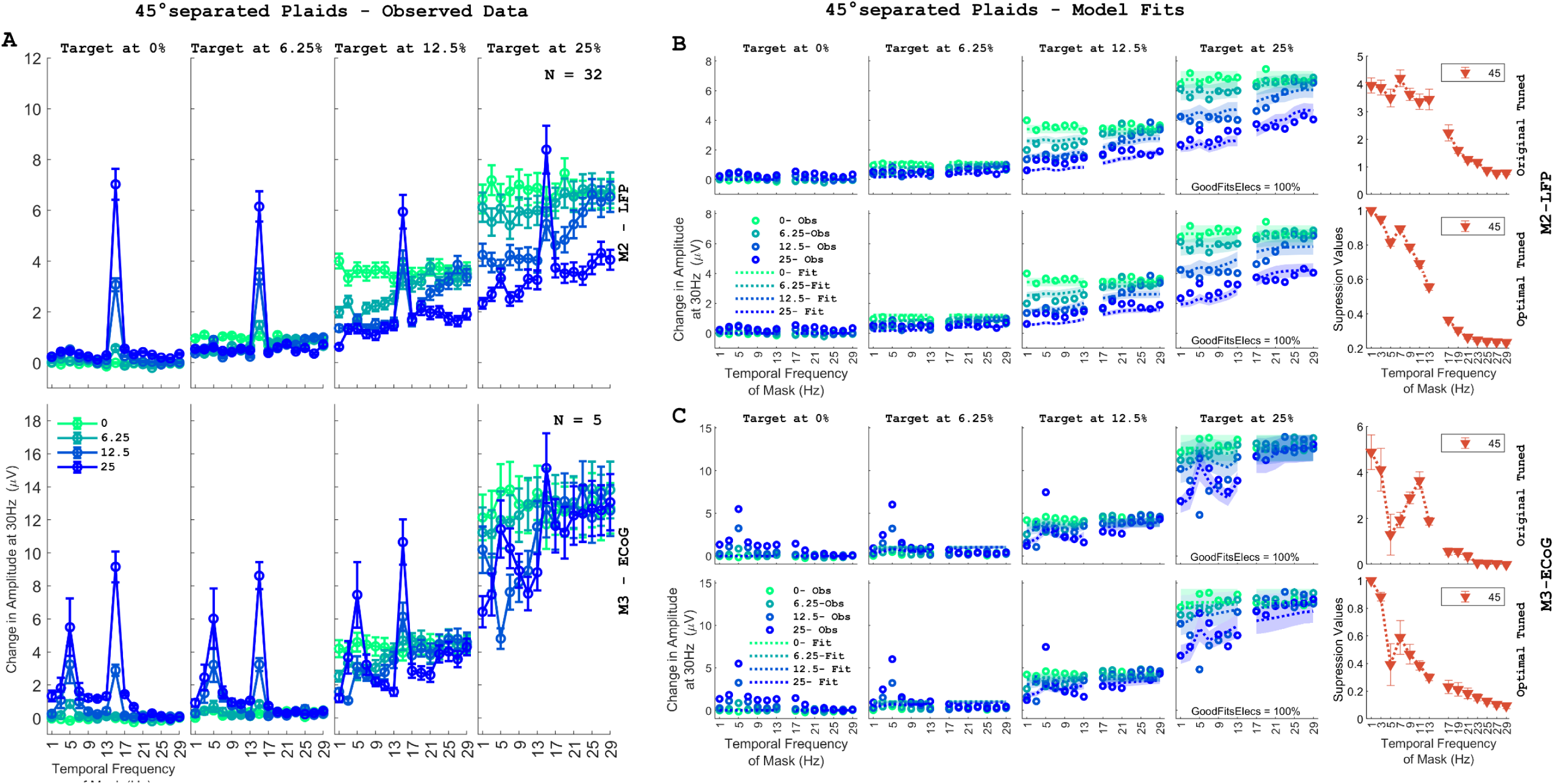
Amplitude response suppression for plaids with 45° orientation difference. (A) Amplitude response at 30Hz averaged across sessions and electrodes for plaid stimuli with 45° separation, with target and mask grating presented at various contrast combinations plotted as a function of temporal frequency of mask grating. Same format as Figure 3. The first row represents the average change in amplitude response from the baseline condition for M2 for a small stimulus. The second row corresponds to the change in amplitude from the baseline for ECoG response in M3 against a full-field stimulus. ‘N’ indicates the number of electrodes for each monkey averaged across sessions. LFP activity was not collected from M1 for these conditions. (B) Empirical and fitted averaged change in amplitude data for electrodes of M2 using original-tuned (upper row) and optimal-tuned normalization model (bottom row), shown in the first four columns. The last column shows the suppression profile obtained from the respective model. The percentage of electrodes having explained variance >= 0.8 is represented as ‘GoodFitsElecs’ in the inset of each row. (C) Same as (B) but for the ECoG grid implanted in M3.

**Figure S3.**
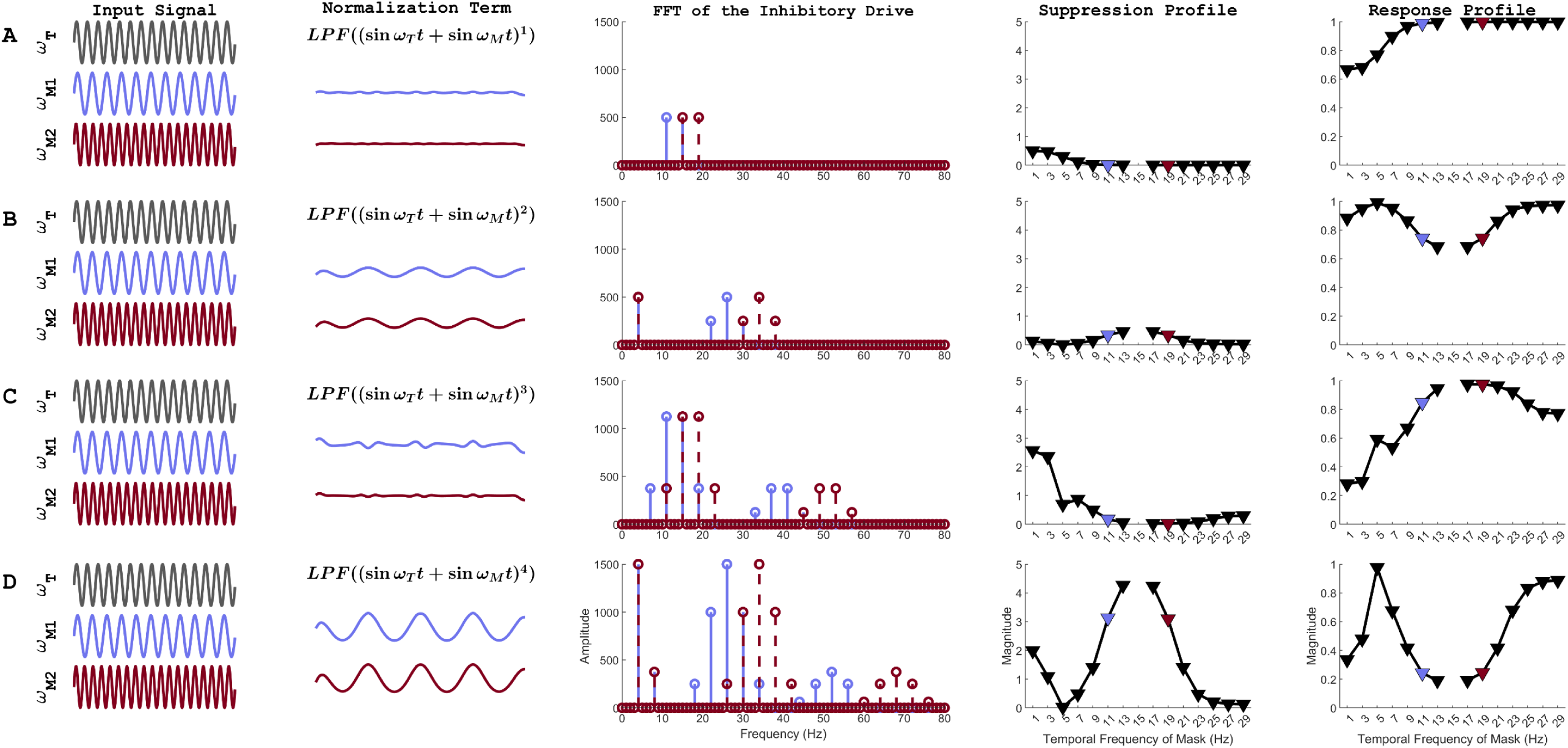
Simulations of low-pass normalization signals with different orders of non-linearity and their respective expected suppression profiles. The five columns correspond to the input signal, inhibitory normalization drive based on summing the sinusoids and then raised to different exponents (exp =1‒ 4 for panels A-D), its Fourier transform, the expected suppression and response profiles. Similar to Figure 5.

**Figure S4.**
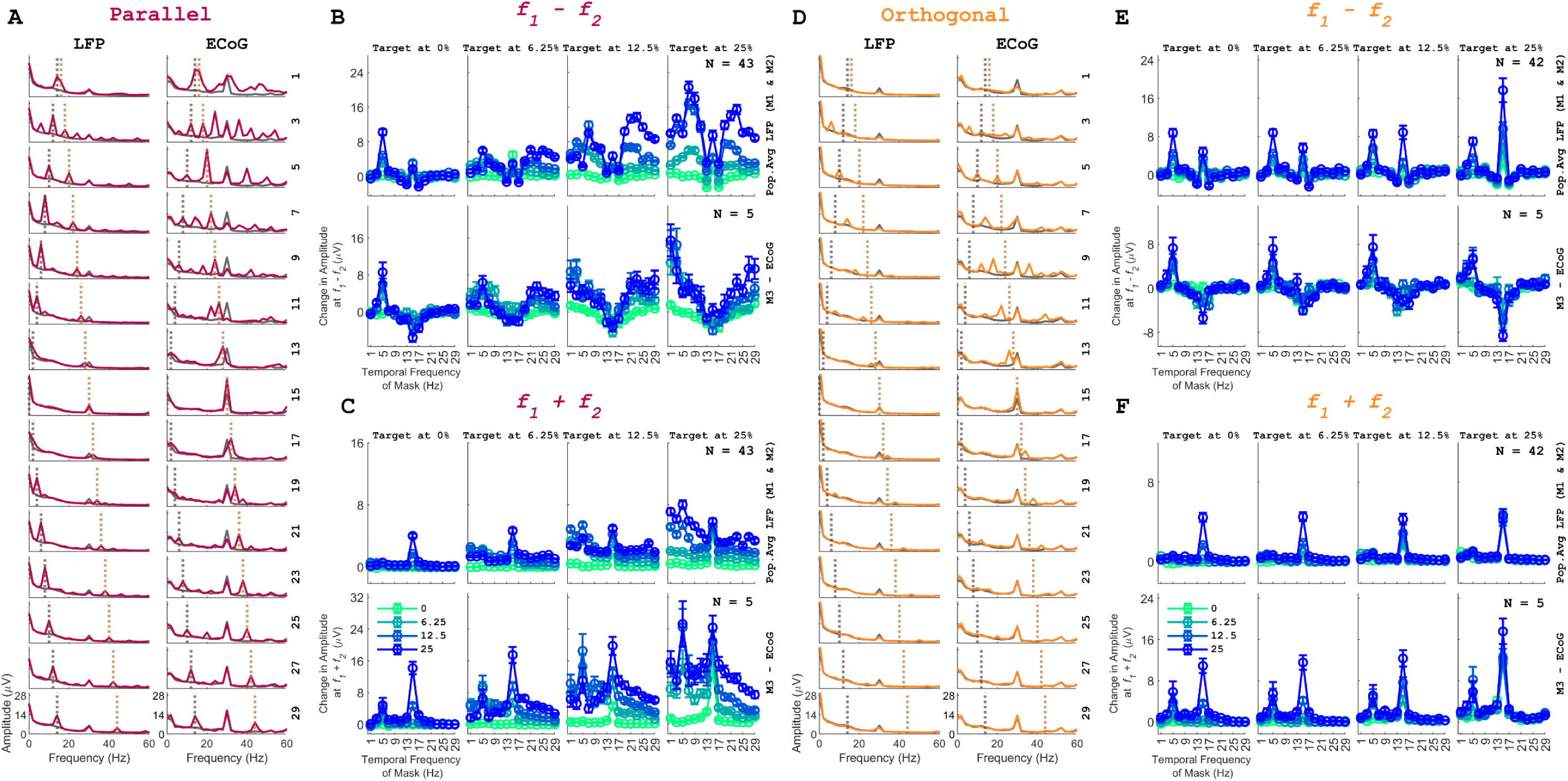
Amplitude response profiles at intermodulation terms for parallel and orthogonal plaids. (A) Mean amplitude spectra averaged across electrodes for parallel plaid sessions (deep magenta trace) with target and mask grating at 25% contrast each for LFP (M1 and M2) and ECoG (M3) activity. Each row corresponds to a mask grating at a different temporal frequency. The grey-coloured trace denotes the grating ‘only’ condition at 25% contrast. The dotted vertical lines indicate intermodulation terms: *f_1_-f_2_* (dark brown trace) and *f_1_+f_2_* (light brown trace). (B) Change in amplitude at plots *f_1_-f_2_* as a function of temporal frequency and contrast of mask grating (green to blue, indicating increasing contrast level) for different contrasts of target grating (different columns). The first row represents the average change in amplitude response from the baseline condition for M1 and M2 for a small stimulus (N = 43). The second row corresponds to the change in amplitude from the baseline for ECoG response in M3 against a full-field stimulus (N=5). (C) Same as (B) but showing change in amplitude at *f_1_+f_2_* instead. The sum-intermodulation components (*f_1_+f_2_*) observed for parallel plaids followed a low-passed profile, with lower mask temporal frequencies evoking a stronger IM component. These results do not align with the suggested model, but they can be obtained when the response itself passes through a low-pass filter. As shown in Figure 8, the SSVEP response in V1 peaks at around 11.51±0.62Hz for small gratings at maximum contrast, so cells are unlikely to respond vigorously to summed IM components with values high as 29+15 = 44 Hz. (D-F) Same as (A-C) but for orthogonal plaid stimuli.

**Figure S5:**
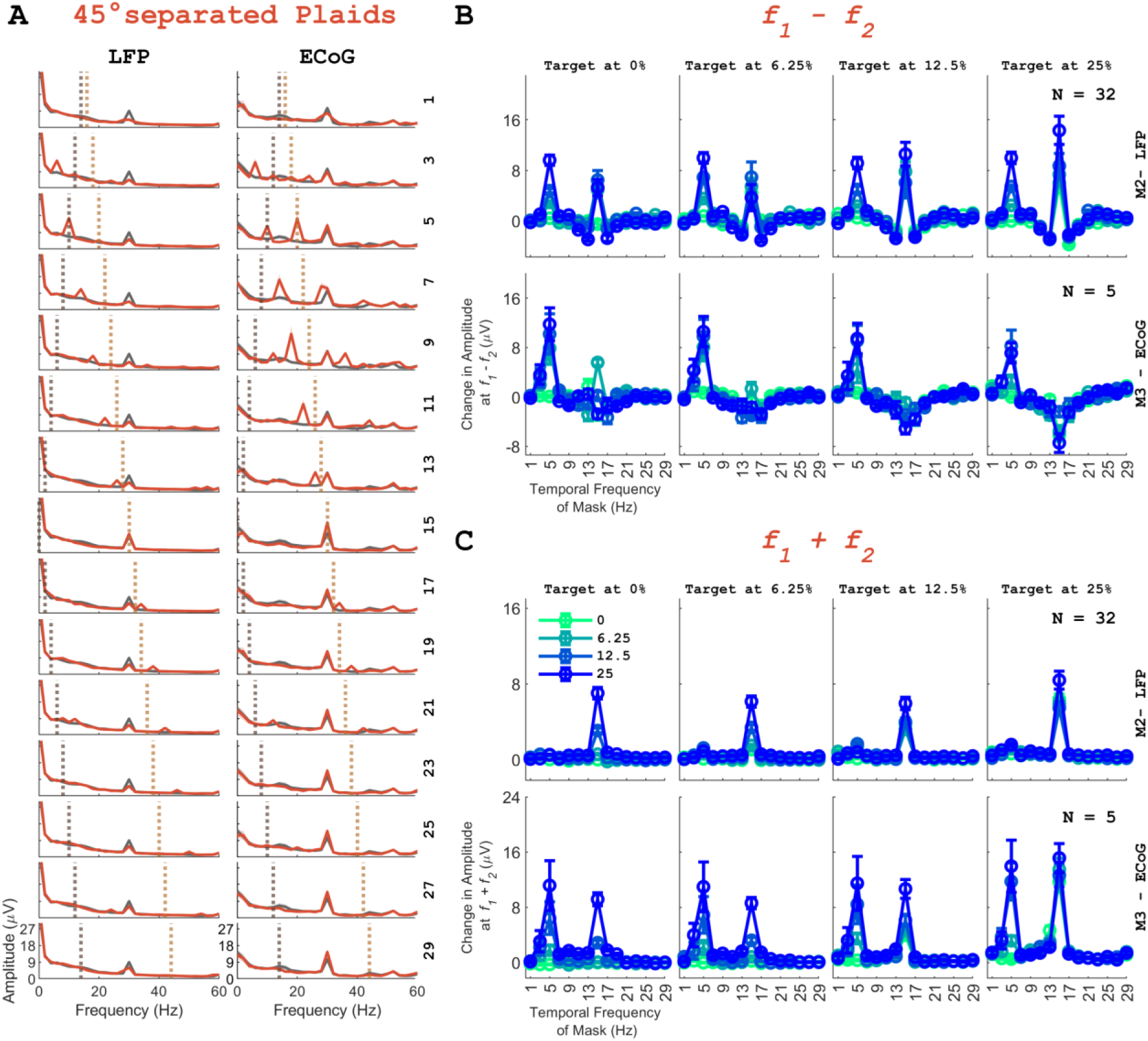
Same as *Figure S4*, panels (A-C), but for 45° separated plaid stimuli presented to M2 and M3. LFP activity was not collected from M1 for these conditions.

**Table S1:**
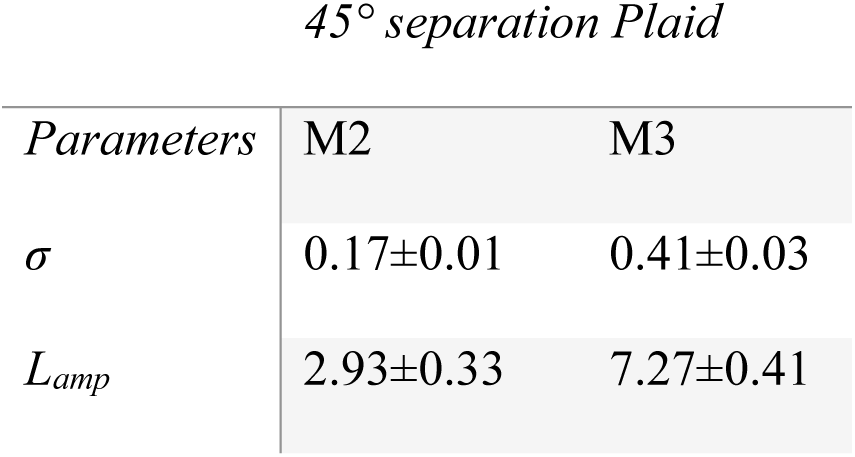
Showing median ± bootstrapped SEM values for parameters obtained from the original-tuned normalization model for 45° separated plaids.

| *Parameters* | M2 | M3 |
| --- | --- | --- |
| $\sigma$ | 0.17±0.01 | 0.41±0.03 |
| $L_{amp}$ | 2.93±0.33 | 7.27±0.41 |

**Table S2:** Showing median ± bootstrapped SEM values for parameters obtained from the optimal-tuned normalization model for 45° separated plaids.

| *Parameters* | M2 | M3 |
| --- | --- | --- |
| $\sigma$ | 0.17±0.01 | 0.42±0.03 |
| $L_{amp}$ | 2.94±0.29 | 7.54±0.25 |
| *Cutoff* | 31.18±0.61 | 23.48±6.13 |
| $\alpha_1$ | 0.16±0.07 | 0.73±0.54 |
| $\alpha_2$ | 4.77±0.40 | 3.90±0.37 |
| $ScF$ | 1±0 | 1±0 |

## REFERENCES

1. Schwartz, O., and Simoncelli, E.P. (2001). Natural signal statistics and sensory gain control. Nat Neurosci 4, 819–825. 10.1038/90526.

2. Ferguson, K.A., and Cardin, J.A. (2020). Mechanisms underlying gain modulation in the cortex. Nat Rev Neurosci 21, 80–92. 10.1038/s41583-019-0253-y.

3. Carandini, M., Heeger, D.J., and Movshon, J.A. (1997). Linearity and Normalization in Simple Cells of the Macaque Primary Visual Cortex. J Neurosci 17, 8621–8644. 10.1523/JNEUROSCI.17-21-08621.1997.

4. Busse, L., Wade, A.R., and Carandini, M. (2009). Representation of concurrent stimuli by population activity in visual cortex. Neuron 64, 931–942. 10.1016/j.neuron.2009.11.004.

5. Cavanaugh, J.R., Bair, W., and Movshon, J.A. (2002). Nature and interaction of signals from the receptive field center and surround in macaque V1 neurons. J Neurophysiol 88, 2530–2546. 10.1152/jn.00692.2001.

6. Tsai, J.J., Wade, A.R., and Norcia, A.M. (2012). Dynamics of normalization underlying masking in human visual cortex. J Neurosci 32, 2783–2789. 10.1523/JNEUROSCI.4485-11.2012.

7. Boynton, G.M., and Foley, J.M. (1999). Temporal sensitivity of human luminance pattern mechanisms determined by masking with temporally modulated stimuli. Vision Research 39, 1641–1656. 10.1016/S0042-6989(98)00199-0.

8. Candy, T.R., Skoczenski, A.M., and Norcia, A.M. (2001). Normalization models applied to orientation masking in the human infant. J Neurosci 21, 4530–4541. 10.1523/JNEUROSCI.21-12-04530.2001.

9. Salelkar, S., and Ray, S. (2020). Interaction between steady-state visually evoked potentials at nearby flicker frequencies. Sci Rep 10, 5344. 10.1038/s41598-020-62180-y.

10. Liza, K., and Ray, S. (2022). Local Interactions between Steady-State Visually Evoked Potentials at Nearby Flickering Frequencies. J Neurosci 42, 3965–3974. 10.1523/JNEUROSCI.0180-22.2022.

11. Burg, M.F., Cadena, S.A., Denfield, G.H., Walker, E.Y., Tolias, A.S., Bethge, M., and Ecker, A.S. (2021). Learning divisive normalization in primary visual cortex. PLoS Comput Biol 17, e1009028. 10.1371/journal.pcbi.1009028.

12. Heeger, D.J. (1992). Normalization of cell responses in cat striate cortex. Vis Neurosci 9, 181–197. 10.1017/s0952523800009640.

13. Carandini, M., and Heeger, D.J. (2012). Normalization as a canonical neural computation. Nat Rev Neurosci 13, 51–62. 10.1038/nrn3136.

14. Reynolds, J.H., and Heeger, D.J. (2009). The normalization model of attention. Neuron 61, 168–185. 10.1016/j.neuron.2009.01.002.

15. Rangel, A., and Clithero, J.A. (2012). Value normalization in decision making: theory and evidence. Curr Opin Neurobiol 22, 970–981. 10.1016/j.conb.2012.07.011.

16. Chen, S., Chen, Y., Geisler, W.S., and Seidemann, E. (2024). Neural correlates of perceptual similarity masking in primate V1. eLife 12, RP89570. 10.7554/eLife.89570.

17. Zhou, J., Benson, N.C., Kay, K., and Winawer, J. (2019). Predicting neuronal dynamics with a delayed gain control model. PLoS Comput Biol 15, e1007484. 10.1371/journal.pcbi.1007484.

18. Mikaelian, S., and Simoncelli, E.P. (2001). Modeling temporal response characteristics of V1 neurons with a dynamic normalization model. Neurocomputing 38–40, 1461–1467. 10.1016/S0925-2312(01)00529-X.

19. Reynaud, A., Masson, G.S., and Chavane, F. (2012). Dynamics of local input normalization result from balanced short- and long-range intracortical interactions in area V1. J Neurosci 32, 12558–12569. 10.1523/JNEUROSCI.1618-12.2012.

20. Fox, R. (1978). Visual Masking. In Perception, S. M. Anstis, J. Atkinson, C. Blakemore, O. Braddick, T. Brandt, F. W. Campbell, S. Coren, J. Dichgans, P. C. Dodwell, P. D. Eimas, et al., eds. (Springer), pp. 629–653. 10.1007/978-3-642-46354-9_20.

21. Regan, D. (1966). Some characteristics of average steady-state and transient responses evoked by modulated light. Electroencephalogr Clin Neurophysiol 20, 238–248. 10.1016/0013-4694(66)90088-5.

22. Norcia, A.M., Appelbaum, L.G., Ales, J.M., Cottereau, B.R., and Rossion, B. (2015). The steady-state visual evoked potential in vision research: A review. Journal of Vision 15, 4. 10.1167/15.6.4.

23. Vialatte, F.-B., Maurice, M., Dauwels, J., and Cichocki, A. (2010). Steady-state visually evoked potentials: focus on essential paradigms and future perspectives. Prog Neurobiol 90, 418–438. 10.1016/j.pneurobio.2009.11.005.

24. Ding, J., Sperling, G., and Srinivasan, R. (2006). Attentional modulation of SSVEP power depends on the network tagged by the flicker frequency. Cereb Cortex 16, 1016–1029. 10.1093/cercor/bhj044.

25. Müller, M.M., Andersen, S., Trujillo, N.J., Valdés-Sosa, P., Malinowski, P., and Hillyard, S.A. (2006). Feature-selective attention enhances color signals in early visual areas of the human brain. Proc Natl Acad Sci U S A 103, 14250–14254. 10.1073/pnas.0606668103.

26. Zemon, V., and Ratliff, F. (1982). Visual evoked potentials: evidence for lateral interactions. Proceedings of the National Academy of Sciences 79, 5723–5726. 10.1073/pnas.79.18.5723.

27. Foley, J.M. (1994). Human luminance pattern-vision mechanisms: masking experiments require a new model. J. Opt. Soc. Am. A, JOSAA 11, 1710–1719. 10.1364/JOSAA.11.001710.

28. Baker, D.H., and Wade, A.R. (2017). Evidence for an Optimal Algorithm Underlying Signal Combination in Human Visual Cortex. Cereb Cortex 27, 254–264. 10.1093/cercor/bhw395.

29. Movshon, J.A., Thompson, I.D., and Tolhurst, D.J. (1978). Receptive field organization of complex cells in the cat’s striate cortex. J Physiol 283, 79–99. 10.1113/jphysiol.1978.sp012489.

30. Adelson, E.H., and Bergen, J.R. (1985). Spatiotemporal energy models for the perception of motion. J Opt Soc Am A 2, 284–299. 10.1364/josaa.2.000284.

31. Adesnik, H., Bruns, W., Taniguchi, H., Huang, Z.J., and Scanziani, M. (2012). A Neural Circuit for Spatial Summation in Visual Cortex. Nature 490, 226–231. 10.1038/nature11526.

32. Li, B., Routh, B.N., Johnston, D., Seidemann, E., and Priebe, N.J. (2020). Voltage-Gated Intrinsic Conductances Shape the Input-Output Relationship of Cortical Neurons in Behaving Primate V1. Neuron 107, 185–196.e4. 10.1016/j.neuron.2020.04.001.

33. Victor, J.D. (1987). The dynamics of the cat retinal X cell centre. J Physiol 386, 219–246. 10.1113/jphysiol.1987.sp016531.

34. Ringach, D.L., Shapley, R.M., and Hawken, M.J. (2002). Orientation Selectivity in Macaque V1: Diversity and Laminar Dependence. J Neurosci 22, 5639–5651. 10.1523/JNEUROSCI.22-13-05639.2002.

35. Meese, T.S., and Holmes, D.J. (2010). Orientation masking and cross-orientation suppression (XOS): Implications for estimates of filter bandwidth. Journal of Vision 10, 9. 10.1167/10.12.9.

36. Petrov, Y., Carandini, M., and McKee, S. (2005). Two Distinct Mechanisms of Suppression in Human Vision. J. Neurosci. 25, 8704–8707. 10.1523/JNEUROSCI.2871-05.2005.

37. Albrecht, D.G., Geisler, W.S., Frazor, R.A., and Crane, A.M. (2002). Visual Cortex Neurons of Monkeys and Cats: Temporal Dynamics of the Contrast Response Function. Journal of Neurophysiology 88, 888–913. 10.1152/jn.2002.88.2.888.

38. Bair, W., Cavanaugh, J.R., and Movshon, J.A. (2003). Time Course and Time-Distance Relationships for Surround Suppression in Macaque V1 Neurons. J. Neurosci. 23, 7690– 7701. 10.1523/JNEUROSCI.23-20-07690.2003.

39. Smith, M.A., Bair, W., and Movshon, J.A. (2006). Dynamics of Suppression in Macaque Primary Visual Cortex. J. Neurosci. 26, 4826–4834. 10.1523/JNEUROSCI.5542-06.2006.

40. Bijanzadeh, M., Nurminen, L., Merlin, S., Clark, A.M., and Angelucci, A. (2018). Distinct Laminar Processing of Local and Global Context in Primate Primary Visual Cortex. Neuron 100, 259–274.e4. 10.1016/j.neuron.2018.08.020.

41. Morrone, M.C., Burr, D.C., and Speed, H.D. (1987). Cross-orientation inhibition in cat is GABA mediated. Exp Brain Res 67, 635–644. 10.1007/BF00247294.

42. Northoff, G., and Mushiake, H. (2020). Why context matters? Divisive normalization and canonical microcircuits in psychiatric disorders. Neuroscience Research 156, 130–140. 10.1016/j.neures.2019.10.002.

43. Ma, W., Liu, B., Li, Y., Josh Huang, Z., Zhang, L.I., and Tao, H.W. (2010). Visual Representations by Cortical Somatostatin Inhibitory Neurons—Selective But with Weak and Delayed Responses. J Neurosci 30, 14371–14379. 10.1523/JNEUROSCI.3248-10.2010.

44. Atallah, B.V., Bruns, W., Carandini, M., and Scanziani, M. (2012). Parvalbumin-Expressing Interneurons Linearly Transform Cortical Responses to Visual Stimuli. Neuron 73, 159–170. 10.1016/j.neuron.2011.12.013.

45. Krishnakumaran, R., Pavuluri, A., and Ray, S. (2025). Delayed Accumulation of Inhibitory Input Explains Gamma Frequency Variation with Changing Contrast in an Inhibition Stabilized Network. J. Neurosci. 45. 10.1523/JNEUROSCI.1279-24.2024.

46. Bertalmío, M., Cyriac, P., Batard, T., Martinez-Garcia, M., and Malo, J. (2017). The Wilson-Cowan model describes Contrast Response and Subjective Distortion. Journal of Vision 17, 657. 10.1167/17.10.657.

47. Malo, J., Esteve-Taboada, J.J., and Bertalmío, M. (2024). Cortical Divisive Normalization from Wilson–Cowan Neural Dynamics. J Nonlinear Sci 34, 35. 10.1007/s00332-023-10009-z.

48. Meese, T.S., and Holmes, D.J. (2002). Adaptation and gain pool summation: alternative models and masking data. Vision Research 42, 1113–1125. 10.1016/S0042-6989(01)00291-7.

49. Holmes, D.J., and Meese, T.S. (2004). Grating and plaid masks indicate linear summation in a contrast gain pool. Journal of Vision 4, 7. 10.1167/4.12.7.

50. Dubey, A., and Ray, S. (2019). Cortical Electrocorticogram (ECoG) Is a Local Signal. J Neurosci 39, 4299–4311. 10.1523/JNEUROSCI.2917-18.2019.

51. Wang, R., Gao, X., and Gao, S. (2004). A study on binocular rivalry based on the steady state VEP. Conf Proc IEEE Eng Med Biol Soc 2006, 259–262. 10.1109/IEMBS.2004.1403141.

52. Silberstein, R.B., Nunez, P.L., Pipingas, A., Harris, P., and Danieli, F. (2001). Steady state visually evoked potential (SSVEP) topography in a graded working memory task. International Journal of Psychophysiology 42, 219–232. 10.1016/S0167-8760(01)00167-2.

53. Zambalde, E.P., Borges, L.R., Jablonski, G., Barros de Almeida, M., and Naves, E.L.M. (2022). Influence of Stimuli Spatial Proximity on a SSVEP-Based BCI Performance. IRBM 43, 621–627. 10.1016/j.irbm.2022.04.003.

54. Li, S., Jin, J., Daly, I., Zuo, C., Wang, X., and Cichocki, A. (2020). Comparison of the ERP-Based BCI Performance Among Chromatic (RGB) Semitransparent Face Patterns. Front Neurosci 14, 54. 10.3389/fnins.2020.00054.

55. Levitt, J.B., Kiper, D.C., and Movshon, J.A. (1994). Receptive fields and functional architecture of macaque V2. Journal of Neurophysiology 71, 2517–2542. 10.1152/jn.1994.71.6.2517.

56. Hawken, M.J., Shapley, R.M., and Grosof, D.H. (1996). Temporal-frequency selectivity in monkey visual cortex. Visual Neuroscience 13, 477–492. 10.1017/S0952523800008154.

57. Hurvich, C.M., and Tsai, C.-L. (1989). Regression and time series model selection in small samples. Biometrika 76, 297–307. 10.1093/biomet/76.2.297.

58. Motulsky, H., and Christopoulos, A. (2004). Fitting Models to Biological Data Using Linear and Nonlinear Regression: A Practical Guide to Curve Fitting (Oxford University Press).

